# Single-cell multi-omic analysis of mesenchymal cells reveals molecular signatures and putative regulators of lung allograft fibrosis

**DOI:** 10.1101/2024.11.27.625698

**Authors:** Lu Lu, A. Patrick McLinden, Natalie M. Walker, Ragini Vittal, Yichen Wang, Stephen T. Russell, Michael P. Combs, Joshua D. Welch, Vibha N. Lama

## Abstract

Survival after lung transplantation is limited by chronic progressive graft failure, termed chronic lung allograft dysfunction (CLAD). Graft-resident mesenchymal cells (MCs) drive CLAD pathogenesis and exhibit stable, dysregulated signaling; however, the transcriptomic and epigenomic drivers behind this fibrogenic transformation remain elusive. Here, we utilize single-cell multi-omic technologies to study gene expression and chromatin accessibility of MCs from the lavage fluid of lung transplant recipients with and without CLAD, obtained either early post-transplantation or after disease onset. MCs obtained after CLAD onset (CLAD-MCs) demonstrated a unique transcriptomic signature compared to non-CLAD controls; a logistic regression model trained on these profiles classified the disease status of individual cells with > 98% accuracy using a set of signature genes. Chromatin accessibility and motif scan analysis identified the CCAAT-enhancer-binding proteins family of transcription factors, specifically CEBPD, as a key marker of the CLAD-enriched subtype. Footprint analysis of early time-point MCs revealed minimal differences in accessibility, suggesting that CEBPD-associated regulatory changes emerge over time after transplantation. Integration and unsupervised clustering identified 8 distinct cell states, and a compositional shift was noted uniquely in CLAD-MCs. Knocking down CEBPD with siRNA in CLAD-MCs partially reverted the CLAD transcriptomic signature, confirming its importance in the dysregulated molecular state of CLAD-associated MCs. scRNA-seq analysis on human lung CLAD tissue provided *in situ* validation of key genes and CEBPD expression changes noted in CLAD-MCs. Our results provide deeper insights into the transcriptomic and epigenomic changes in post-transplant MCs, nominating biomarkers and disease-associated factors with implications for future therapeutic efforts.

## Introduction

Mesenchymal cell (MC) infiltration and matrix deposition leading to architectural distortion are key features of fibrotic diseases across all organs (1). Significant insight has been gained into the mechanisms of fibrosis by *in vitro* investigations of MCs derived from diseased human tissues. While initial focus was on mediators present in the pro-fibrotic milieu that induce and sustain myofibroblast differentiation (2, 3), a novel paradigm of an autonomous fibrotic transformation of MCs has recently emerged with demonstration of a myriad of stable changes at the transcriptional and translational level in these MCs removed from their microenvironment (4–8). Recent advancements in single-cell sequencing technologies provide an unprecedented opportunity to improve upon this work by delving into the intricate heterogeneity of cells in culture and studying genetic and epigenetic molecular signatures in an unbiased manner.

Lung transplantation, the only viable option available to patients with end-stage lung diseases, presents a formidable challenge with a median survival of 5.8 years, the lowest among all solid organ transplants (9). The major cause of these poor long-term outcomes is chronic lung allograft dysfunction (CLAD), characterized by recalcitrant fibrotic remodeling of the allograft and progressive lung function decline (10). CLAD develops in greater than 50% of patients by 5 years of transplantation and is the major cause of mortality after 1-year post-lung transplantation (10). Allograft fibrosis, emanating from the bronchovascular bundles and small airway fibrosis or bronchiolitis obliterans, is a key feature of the failing grafts in CLAD. Therapeutic approaches to modulate allograft fibrogenesis are a key unmet need in this arena, as augmented immunosuppression fails to alter the progression of chronic graft failure meaningfully.

A significant body of translational work from our laboratory has highlighted a stable alteration in graft-resident MCs that perpetuates fibrosis in a chronically rejecting lung allograft. We were the first to discover a resident population of mesenchymal progenitors in human adult lungs by demonstrating that MCs cells isolated from bronchoalveolar lavage (BAL) of human lung allografts are of donor origin and retain the expression of embryonic lung mesenchyme-associated transcription factor Forkhead box gene *FOXF1* (11, 12). These transcriptionally distinct *Foxf1*^+^ subset of collagen I-expressing lung-resident MCs were shown to reside topographically along the bronchovascular bundles and contribute to allograft fibrosis (11, 13). Translational human studies demonstrated that their expansion (14, 15) and fibrotic differentiation (6, 7, 16–18) precede and accompany the development of CLAD. Mechanistic investigations have revealed stable dysregulation across a myriad of signaling pathways in human CLAD lung-derived MCs, suggesting a yet undiscovered alteration in global transcriptional and epigenomic landscape (6, 7, 11, 12, 16–18). Whether the altered fibrogenic state of these graft-resident MCs is acquired over time after transplantation also remains to be investigated.

In this study, we leverage single-cell multi-omics profiling and provide an unbiased examination of distinct MC gene expression and chromatin accessibility signatures in CLAD. Our findings elucidate the molecular signatures associated with CLAD and highlight temporal changes in human lung allograft-derived MCs marked by unique gene expression profiles, regulatory networks, and epigenetic modifications. We nominate putative regulators of the dysregulated molecular states of CLAD MCs and specifically highlight a central role for transcription factor (TF) CCAAT/enhancer binding protein delta (CEBPD) in sustaining the fibrotic state of MCs. Additional single-cell RNA sequencing (scRNA-seq) analysis of MCs from human CLAD and normal lungs further demonstrated that cultured MCs retain and reflect key transcriptomic signatures from their *in vivo* counterparts.

## Results

### Single-cell multi-omic profiling of human mesenchymal cells from lung transplant recipients

To investigate if CLAD is associated with stable transcriptomic and epigenomic changes in graft MCs, joint scRNA-seq and single-cell ATAC sequencing (scATAC-seq) was performed on MCs isolated from BAL of lung transplant recipients (**Figure 1; Additional file 2: Table S1**). MCs from patients who had a diagnosis of CLAD at the time of BAL (CLAD-MCs) were compared to the MCs obtained from control CLAD-free patients (non-CLAD MCs). The incidence of CLAD increases with time post-transplant, and CLAD diagnosis is generally made after the first year of lung transplantation. To investigate if transcriptional and epigenomic changes are inherent or acquired, MCs were also studied from BAL samples obtained early in the course (<6 weeks post-transplant). These patients were later clinically classified as having developed CLAD or remained CLAD-free. Single-cell multiome sequencing with the 10X Genomics Chromium platform was performed. To lower library preparation costs, MCs were pooled before loading them onto the 10X Chromium, allowing profiling of multiple donors simultaneously. Each donor was genotyped using an Illumina SNP array. scRNA-seq and scATAC-seq reads were demultiplexed using this genotype information (**Figure 1**, see Methods). After quality control filtering, we obtained 22,918 MCs in total for further analysis (**Additional file 1: Figure S1**).

**Figure 1.**
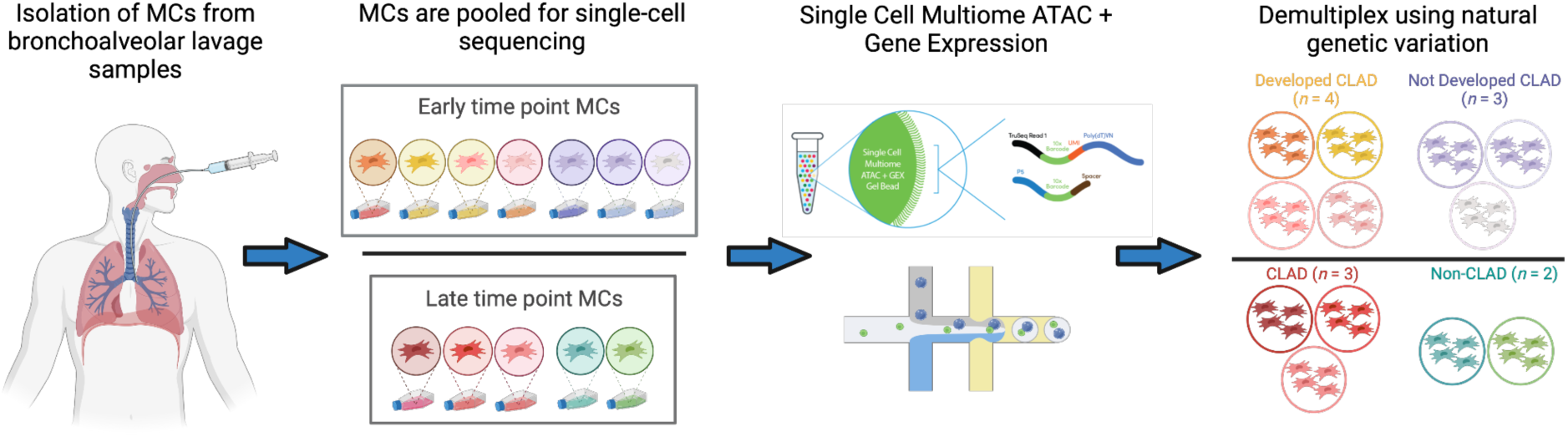
**Experimental strategy**. Mesenchymal cells (MCs) were isolated from bronchoalveolar lavage (BAL) fluid derived from lung transplant recipients. MCs were screened using clinical parameters and then pooled together prior to single-cell multiome profiling. The donor identity of each cell was determined using genetic variants captured in the gene expression and chromatin accessibility reads.

### Classification models predict CLAD status of patient-derived MCs from single-cell gene expression

To identify transcriptomic markers of CLAD in patient-derived MCs, we trained classification models to distinguish cells originating from CLAD versus non-CLAD lungs (**Figure 2A**). To determine the optimal number of genes in the classifier, we quantified the model performance while varying the parameter α, which controls the number of genes with nonzero regression coefficients. Higher values of α indicate a stronger regularization and result in fewer genes selected. The accuracy score and the corresponding number of genes for each α-value were reported (**Figure 2B**). The classifiers achieved high accuracy, with the highest test accuracy of 99.2% on classifying the late-time-point MCs (**Figure 2B**). In addition, we calculated the Jaccard similarity index between each set of selected genes across different parameter values (**Figure 2C**). The high similarity score between adjacent pairs of α-values indicates that a key set of genes persists across a range of α parameters. To ensure that our model learned the true biological signal of CLAD status, rather than simply biological variance among donors, we additionally evaluated our model by holding out all cells from one donor at a time. We observed similar performance training on all donors compared to one donor leave-out, indicating that the model is not simply capturing biological variation among donors (**Additional file 1: Figure S2**).

**Figure 2.**
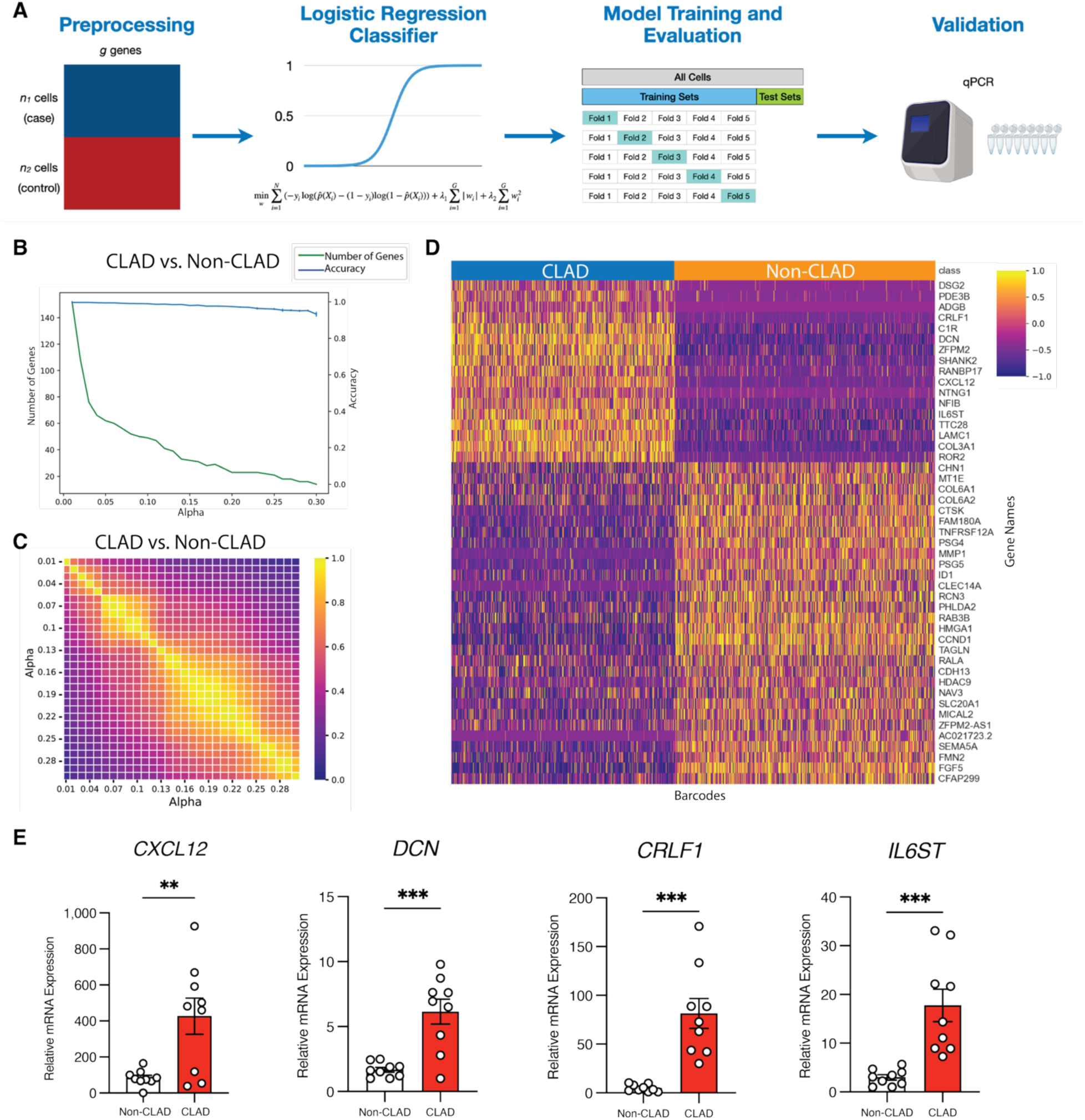
Classifying CLAD status from single-cell gene expression. **(A)** Workflow of training logistic regression classifier. **(B)** The blue lines indicate average accuracy score of the logistic regression classifiers from 5-fold cross-validation. Error bars indicate the variance in accuracy across folds. The green lines indicate the number of genes with non-zero coefficients from the logistic regression classifiers trained on the test sets. **(C)** Heat-map showing the overlap of genes among different coefficients. **(D)** Heatmaps depicting the expression levels of gene signatures in CLAD patient-derived MCs constructed from the logistic regression classifiers with α of 0.1. Each column represents a cell, and each row represents one gene colored by gene expression level. **(E)** RT-qPCR of *IL6ST*, *CRLF1*, *CXCL12*, and *DCN* expression from CLAD and non-CLAD cell lines. (*p<0.05; **p ≤ 0.01)

When applying the same classification approach to the early time-point MCs, we found that classification was more challenging compared to the late time-point MCs. The highest test accuracy achieved was 91.2%, and the classifier required more gene features to attain this performance (**Additional file 1: Figure S3A**). This suggests that there is less transcriptional difference among the early time-point MCs than the late time-point. Differential expression analysis further supports this observation that only 66 genes were significantly differentially expressed in the early time-point MCs. In comparison, 1,707 genes were significantly differentially expressed in the late time-point MCs. (**Additional file 1: Figure S3B**)

To select a final α value for subsequent analyses, we identified an “elbow” in the plot of accuracy vs. number of genes, choosing a model that uses as few genes as possible without a significant drop in accuracy. We opted for an α of 0.1, which identified 47 genes for the late time-point group (**Figure 2D**), and an α of 0.05, resulting in the selection of 54 genes for the early time-point group (**Additional file 1: Figure S3C**). These classifiers achieved test accuracy of 98.7% and 85.8% in the late and early time-point groups, respectively. Many of the genes selected by these classifiers are plausibly linked to CLAD based on previous studies. The selected genes for CLAD include *DCN*, a known gene involved in fibrogenesis (19). We observed that *CXCL12*, an important homeostatic chemokine implicated in both homeostatic and inflammatory contexts (20–24), was upregulated in CLAD-MCs (**Figure 2D**). To verify differential expression of select genes discovered through scRNA-seq of CLAD-MCs, we performed RT-qPCR for *CXCL12, DCN, CRLF1, IL6ST* in MCs isolated from additional CLAD and non-CLAD donors (**Figure 2E**). We successfully verified increased mRNA expression in CLAD-MCs for each of the four genes examined (**Figure 2E**).

### Chromatin accessibility analysis highlights CEBPD as a key positive regulator in CLAD patient-derived MCs

We next examined differences in chromatin accessibility profiles between CLAD-and non-CLAD-MCs. First, we performed peak calling on each sample, then merged the peaks, identifying 114,464 accessible peaks that were concordant among donors. Quality control to filter cells was performed for each sample separately (See Methods; **Additional file 1: Figure S4**). To find peaks with differential accessibility in each condition, we performed Wilcoxon rank-sum tests on CLAD and non-CLAD samples. Among 114,464 accessible peaks, we identified 2,983 (2.61%) up-regulated peaks and 4,120 (3.60%) down-regulated peaks in CLAD by setting filtering thresholds of FDR ≤ 0.05 & abs(Log2FC) ≥ 1 (**Figure 3A**).

**Figure 3.**
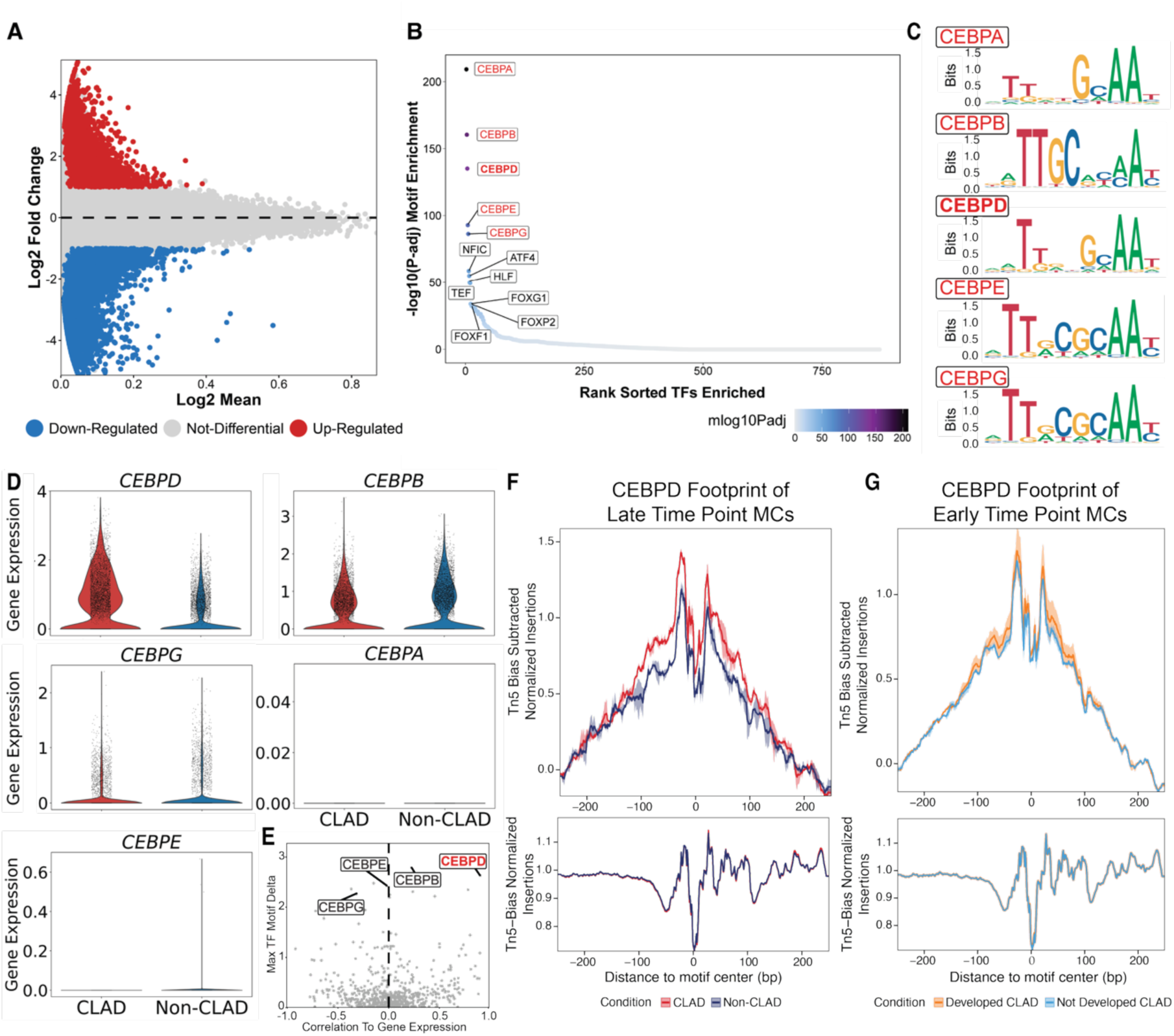
Single-cell chromatin accessibility analysis to identify epigenetic markers of CLAD MCs. **(A)** MA plot showing marker peaks in CLAD. **(B)** TF motifs enriched in CLAD ranked by enrichment. **(C)** Sequence logo of motifs from CEBP family (CEBPA, CEBPB, CEBPD, CEBPE, and CEBPG) in position weight matrices (PWMs). **(D)** Violin plots depicting the expression levels of CEBP family (CEBPA, CEBPB, CEBPD, CEBPE, and CEBPG) across conditions. The width of each plot represents the distribution of expression values. Comparisons are shown between late-time-point MCs with (red) or without (blue) CLAD. **(E)** Dot plot showing correlation between motif accessibility and its corresponding TF expression. The x-axis is Pearson correlation coefficient between motif deviation z-scores from chromVAR and its corresponding TF expression. The y-axis is the TF deviation z-scores from chromVAR. TFs from CEBP family are annotated and CEBPD is highlighted in red. **(F-G)** CEBPD footprinting of late-time-point **(F)** and early-time-point **(G)** MCs.

To understand the key transcription factors (TFs) that could bind to these accessible chromatin sites, we then performed a motif scan. The CEBP family of transcription factors, including CEBPD, a TF involved in proliferation, apoptosis, and possibly fibrogenesis (25–27), was noted to be enriched in CLAD (**Figure 3B**). However, members of CEBP family share high similarities in their binding motifs, which can be visualized as sequence logos derived from position weight matrices (PWMs) (**Figure 3C**) (28). This motif similarity is common among many TF families, making it challenging to pinpoint the specific TFs that might be involved in regulation (29). To address this, we incorporated the matched gene expression data. We first examined the expression of the CEBP family and found that CEBPD is the only TF among the CEBP family that is enriched in CLAD. It has 32-fold higher expression in CLAD-MCs than in non-CLAD MCs (**Figure 3D**). CEBPB showed similar expression levels between the two conditions, while CEBPA, CEBPE, and CEBPG were either lowly expressed or not expressed at all (**Figure 3D**). We further calculate a Pearson correlation between the gene expression of TFs and the chromatin accessibility of their corresponding motif. CEBPD yielded the highest correlation of 0.9 (adj-p = 2.43e-179) (**Figure 3E**). Moreover, we performed footprinting analysis at predicted CEBPD binding sites. When a TF is bound to DNA, it physically protects that region from transposase activity during ATAC-seq, creating a characteristic footprint: a localized dip in accessibility at the center of the motif, flanked by regions of higher accessibility. This footprint pattern was clearly observed at CEBPD motifs, indicating active binding of CEBPD (**Figure 3F**). The footprint pattern exists in both conditions, but the significant difference in accessibility suggests a change in binding affinity between CLAD and non-CLAD MCs, which may drive the gene regulatory differences. In contrast, footprint analysis of early time-point MCs revealed minimal differences in accessibility, suggesting that CEBPD-associated regulatory changes emerge later in the disease course and are not present shortly after transplantation (**Figure 3G**). Taken together, these results point to CEBPD as a candidate TF driving gene regulatory alterations in CLAD MCs and such alterations do not take place early in the transplantation.

### Single-cell analysis reveals temporal cell state shifts in patient-derived MCs

To explore the compositional heterogeneity in MCs derived from lung transplant patients across different time points, we performed integration and unsupervised clustering using LIGER (30, 31). We integrated samples from 12 donors to identify shared and distinct cell states between CLAD and control cells despite biological variation among donors (**Figure 4, A and B**). We then annotated clusters using top Gene Ontology (GO) terms from differentially expressed genes and enriched TF motifs (**Figure 4A**). We also compared published marker genes from previously identified fibroblast cell states, and these published studies were utilized to help with the nomenclature (32, 33). Eight distinct cell states were identified and annotated as proliferating, nerve-associated, immune-interacting, vascular-interacting, intermediate, pre-fibrotic, fibrotic, and inflammatory (**Figure 4A**). The marker genes and GO terms for the vascular-interacting and immune-interacting populations are concordant with those identified in a previous cross-tissue study of fibroblast cell states (32). Immune-interacting clusters are enriched for GO terms related to humoral immune response (34) and cytokine-mediated signaling. The nerve-associated cluster exhibits GO terms associated with neurogenesis and neuronal development. The features present in this cluster are similar to recently discovered fibroblasts related to airway peripheral nerves (33). The proliferating cell state is distinguished largely by expression of mitotic cell cycle genes, microtubule and cytoskeleton genes, and other markers of proliferation. GO terms enriched in the fibrotic cluster included positive regulation of transcription by RNA Polymerase II, cell surface receptor signaling pathway, regulation of cell differentiation, cell communication, signal transduction, and cell motility. These included genes such as *CRLF1*, *VEGFA*, *HDAC4*, *ITGB8* and *CXCL12*. Molecular processes identified by GO analysis in this cluster included terms such as CD4 receptor binding and extracellular matrix structural constituents with genes such as *IL16*, *PLSCR4*, *PLSCR1*, various collagens, *MRXA5,* and *LAMB1*. The inflammatory cluster shows upregulation of neutrophil-mediated immunity and neutrophil activation involved in immune responses.

**Figure 4.**
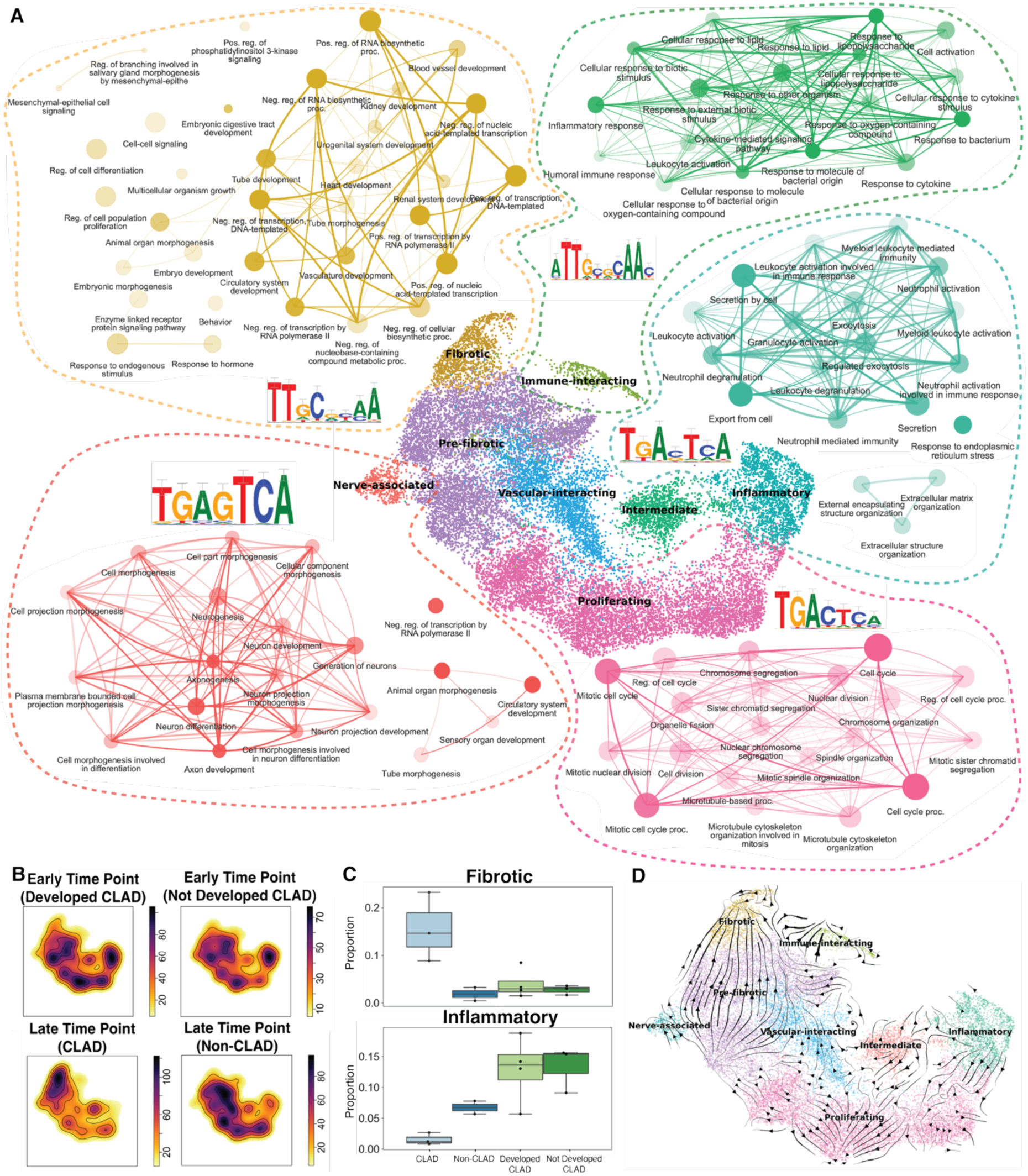
Cell state heterogeneity and cell state shifts in patient-derived MCs. **a** Uniform manifold approximation and projection (UMAP) representation of 22,918 cells derived from lung transplant recipients at the two time points. Each dot represents a single cell, and cells are colored by clusters. **(B)** Density plots of integrated MCs (top left: early time-point (later developed CLAD), top right: early time-point (later not developed CLAD), bottom left: late time-point (CLAD patient-derived), bottom right: late time-point (Non-CLAD patient-derived). **(C)** Compositional change in fibrotic (top) and inflammatory (bottom) clusters. **(D)** Velocities fit by VeloVAE are visualized as streamlines in the UMAP plot.

Having characterized the cell state diversity of the MCs, we next looked for cell state shifts/compositional differences between MCs across different conditions (**Figure 4B**). A striking observation was the unique pattern that emerged for CLAD-MCs where there was a shift towards fibrotic state. Most of the cells in the fibrotic cluster came from CLAD patients (**Figure 4C**). The inflammatory state was predominantly enriched in early time-point MCs, possibly reflecting innate immune engagement during the early post-transplant period. Notably, the composition of early time-point MCs was highly similar between patients who later developed CLAD and those who did not develop CLAD, with MCs collected from patients who later developed CLAD having a slight shift towards the fibrotic state. These findings are consistent with previous observations that transcriptional and epigenetic differences between CLAD and non-CLAD MCs emerge later in the disease course and are not apparent shortly after transplantation. Collectively, these results indicate that MCs from different time points and conditions occupy different parts of the fibroblast cell state space.

RNA velocity analysis using VeloVAE (35) to investigate potential relationships among the cell states was performed (**Figure 4D**). The primary cellular state transition suggested was from pre-fibrotic to fibrotic, with no clear unidirectional relationship between other states.

### Enhancer-driven gene regulatory network analysis reveals putative regulators of the CLAD-enriched fibrotic state

The joint gene expression and chromatin accessibility data we collected provided a unique opportunity to investigate regulatory activity involving the expression of TFs, the chromatin accessibility of TF-binding sites, and the expression of target genes within MCs among different cellular states. For this purpose, we used SCENIC+ to infer an enhancer-driven gene regulatory network (36). SCENIC+ looks for three-way relationships in which TFs are expressed in cells where peaks containing binding sites for the TFs are accessible, and the accessibility of the binding sites is also correlated with the expression of nearby genes. In total, this analysis identified 447 TFs that are predicted to regulate downstream target genes. Among these TFs, we identified 24 activator eRegulons (correlation coefficient > 0.5) predicted to target a total of 12,174 regions and 3,703 target genes. We also calculated the cell state specificity of each activating TF (**Figure 5A**). For example, one notable activating factor is NFIB, a TF known to be expressed in lung MCs (37). NFIB shows the highest activity in the nerve-associated state (**Figure 5A**). Research has demonstrated that NFIB plays an important role in the development and progression of various nerve-related diseases (38, 39). Moreover, HMGA1, a regulator enriched in the inflammatory cluster, has been shown to be involved in inflammatory pathways (40). This is consistent with our observation that, early after lung transplantation, MCs present an inflammatory state. CEBPD, the transcription factor found to be enriched in the CLAD state, showed the highest activity in the fibrotic cluster.

**Figure 5.**
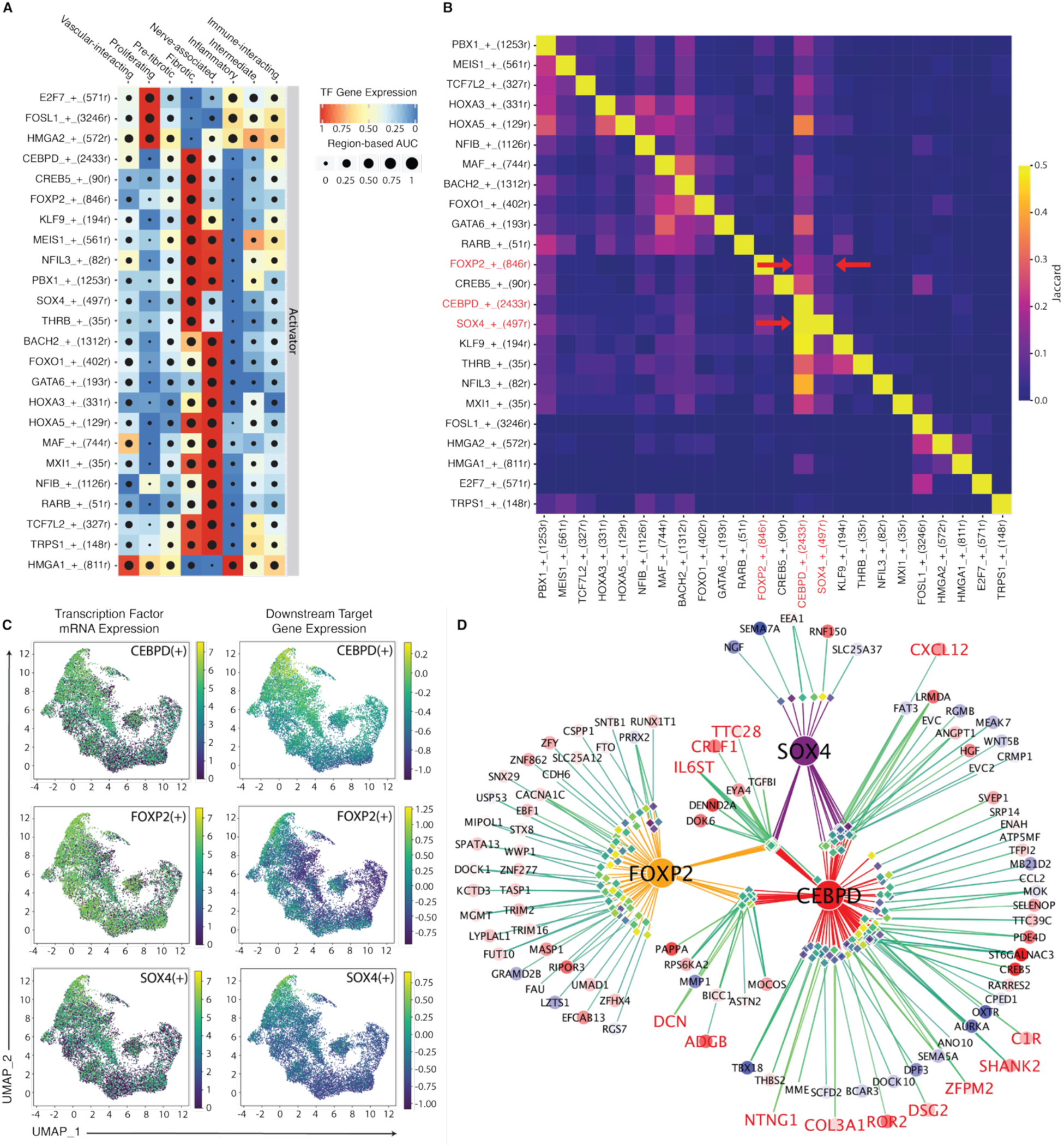
Gene regulatory analysis combining single-cell trancriptomic and epigenomic features nominates regulators of CLAD-enriched fibrotic state. **(A)** Dot plot showing the state-specificity of the predicted targets for each TF. Each row is a cell state and each column is a TF. The color of each square indicates the aggregate expression of predicted target genes for each TF in each cell state. Dot size indicates the area under the recovery curve (AUC) for a classifier trained to predict the cell state from the chromatin accessibility of target peaks. Higher AUC means that the target regions for the TF are more state-specific. **(B)** Heatmap showing overlap between predicted target regions of each TF. Each square shows the Jaccard similarity index (intersection of target regions divided by the union of target regions) for the pair of TFs in the corresponding row and column. Higher values indicate more overlap between the targets of the pair of TFs. **(C)** UMAP visualization of CEBPD, FOXP2, and SOX4 and predicted targets. Each dot represents a cell. Dots are colored by TF expression (left) or the sum of target gene expression (right). **(D)** Visualization of the enhancer-driven gene regulatory network formed by CEBPD, FOXP2, and SOX4. Each circle is a gene; color and size represent the logFC of the gene’s expression in fibrotic MCs compared to the rest of the cells. Thus, a large, red circle indicates a gene with much higher expression in fibrotic MCs and a large blue circle indicates a gene with much lower expression in fibrotic MCs. Each diamond is a chromatin accessibility peak (putative enhancer) and its color represents the logFC of the accessibility in fibrotic MCs compared to the rest of the cells (yellow is higher accessibility in fibrotic MCs). Each line indicates a regulatory relationship; line color indicates the strength of the correlation between peak and gene (higher correlation is more yellow). Gene symbols in red letters are part of the CLAD signature from Figure 2E.

We further examined activating factors enriched in the fibrotic cell state. The factors with the most state-specific expression in the fibrotic state and the most predicted targets are CEBPD (2,433 targets), FOXP2 (846 targets), and SOX4 (497 targets). Although PBX1 (1,253 targets) and MEIS1 (561 targets) also have high expression in the fibrotic state and a significant number of predicted targets, they additionally show significant activity in the nerve-associated state, indicating that their activity is less specific to the fibrotic state. Interestingly, the predicted targets of CEBPD, FOXP2, and SOX4 substantially overlap, while PBX1 and MEIS1 are predicted to regulate a separate set of genes that overlap between the two factors but are largely disjoint from genes regulated by CEBPD, FOXP2, and SOX4 (**Figure 5B**). The aggregate expression of predicted gene targets of CEBPD, FOXP2, and SOX4 shows strong enrichment in the fibrotic state (**Figure 5C**). Strikingly, this aggregate expression is more state-specific than the gene expression of the TFs themselves, likely reflecting state-specific differences in the chromatin accessibility of the TF binding motifs.

We further investigated the predicted gene and peak targets of CEBPD, FOXP2, and SOX4. We plotted their gene regulatory network to confirm the observation of cooperativity among and visualize the target genes of the factors (**Figure 5D**). Interestingly, 13 of these target genes were also predicted by our logistic regression model as key up-regulated genes related to CLAD (**Figure 2E**). Several of the CLAD signature genes, including *DCN*, *CXCL12*, *IL6ST*, *CRLF1*, *TTC28*, and *ADGB* are also predicted to be co-regulated by 2 or more of the factors.

### CEBPD knockdown partially reverts fibrotic gene expression signatures

Given that CEBPD is noted to be enriched in CLAD disease state and is the fibrotic cluster-specific factor with the most predicted targets (and the most predicted targets that are also part of our CLAD signature), we decided to further explore the regulatory role of CEBPD. To do this, we knocked down CEBPD using small interfering RNAs (siRNAs). CLAD-MCs were treated with either a CEBPD-targeting siRNA (treated group) or a scrambled siRNA (control group) for 16 hours. Knockdown efficacy was confirmed at both the mRNA and protein levels using qPCR and Western blots (**Figure 6, A and B**). To measure the gene expression changes induced by CEBPD silencing, we performed bulk RNA-seq on both the treated and control groups across the four CLAD samples. After sequencing, we obtained more than 36 million reads per sample mapped to genes. Differential expression analysis revealed a total of 2,436 differentially expressed genes between control and knockdown samples, with 1,265 genes upregulated and 1,171 genes downregulated after CEBPD silencing (**Figure 6C**). The overlap between differentially expressed genes from the bulk RNA-seq and the predicted downstream targets of CEBPD from the eGRN was examined. Of the 593 genes predicted to be downstream targets of CEBPD as an activator, 85 were indeed downregulated after CEBPD silencing, while 64 remained upregulated (**Additional file 1: Figure S6A**). Additionally, of the 201 genes predicted to be downstream targets of CEBPD as a repressor, 11 were upregulated and 32 remained downregulated. Next, we compared the differentially expressed genes from the bulk RNA-seq with the previously identified CLAD signature in Figure 2 from the logistic regression model (**Additional file 1: Figure S6B**). None of the upregulated genes in CLAD showed significantly higher expression in the CEBPD-silenced group than in the control group. Notably, 5 of the 17 upregulated genes in CLAD, including *CXCL12* and *IL6ST*, were significantly downregulated in the bulk RNA-seq (**Figure 6C**).

**Figure 6.**
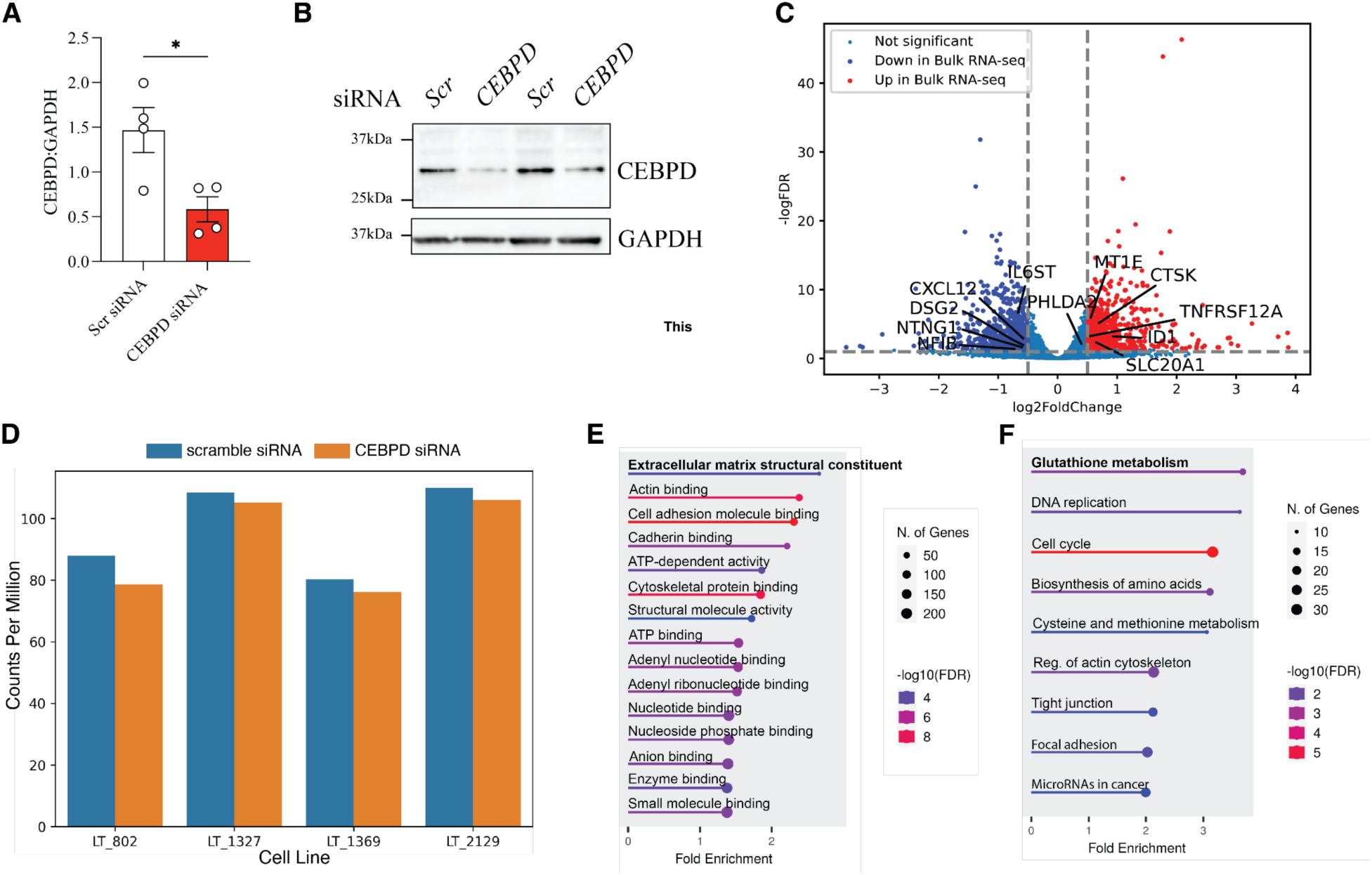
CEBPD knockdown partially reverts CLAD gene expression signature. **(A)** RT-qPCR expression of *CEBPD* relative to GAPDH in CLAD MCs treated with scrambled or *CEBPD* siRNA (*p<0.05). **(B)** Protein expression of CEBPD in CLAD MCs treated with scrambled or CEBPD siRNA as measured by Western blot analysis. **(C)** Volcano plot illustrating the differentially expressed genes following *CEBPD* knockdown in CLAD MCs. The x-axis represents the log2 fold change, and the y-axis represents the-logFDR. Genes that are significantly upregulated (log2FC > 0.5, adj. p-value < 0.05) are marked in red, while significantly downregulated genes (log2FC <-0.5, adj. p-value < 0.05) are marked in blue. Non-significant genes are shown in gray. Gene symbols highlighted with texts are part of the CLAD signature from Figure 2E. **(D)** Bar plot showing the total expression of the upregulated CLAD signature genes between CLAD MCs treated with scrambled (blue) or *CEBPD* (orange) siRNA. The x-axis is the cell line; control and knockdown samples from the same line are shown as adjacent bars. The y-axis represents the log sum of counts per million. **(E)** GO enrichment analysis showing downregulation of ECM-related genes after *CEBPD* knockdown. **(F)** KEGG pathway enrichment analysis showing downregulation of glutathione metabolism-related genes after *CEBPD* knockdown.

Similarly, 6 out of 30 downregulated genes in CLAD were significantly upregulated in the bulk RNA-seq after knockdown (**Figure 6C**). Furthermore, we added up the total expression of all 17 upregulated signature genes in both control and knockdown cells and found that the total expression of the upregulated CLAD signature genes (in units of counts per million) was lower in each of the four knockdown samples compared to its donor-matched control sample (**Figure 6D**). These results suggest that silencing CEBPD partially reverts the CLAD gene expression signature we identified.

To further understand the transcriptomic changes induced by CEBPD knockdown, we performed GO enrichment analysis and examined enriched KEGG pathways (**Figure 6 E and F**). The top GO terms identified as downregulated were extracellular matrix (ECM) structural constituent and actin binding, with decreased expression of genes such as *COL11A1*, *COL12A1*, *FBN2*, and *POSTN*. KEGG pathway also identified downregulation in important pathways of glutathione metabolism, DNA replication, cell signaling, and biosynthesis of amino acids. These data suggest an important role for CEBPD in regulating MC activation and fibrotic differentiation.

### Cultured MCs retain and reflect their *in vivo* transcriptomic signatures

To determine whether our cultured MCs retain and reflect their *in vivo* phenotype, we analyzed MCs directly isolated from CLAD human lung autopsy samples. We collected and performed scRNA-seq on cells from human lung autopsy samples, including one from a CLAD patient and two from healthy controls. Due to the low number of MCs detected in these initial samples (**Additional file 1: Figure S7A**), we sequenced two additional samples (one from the same CLAD patient and one from the same healthy control) after CD45^+^ depletion (**Figure 7A**).

**Figure 7.**
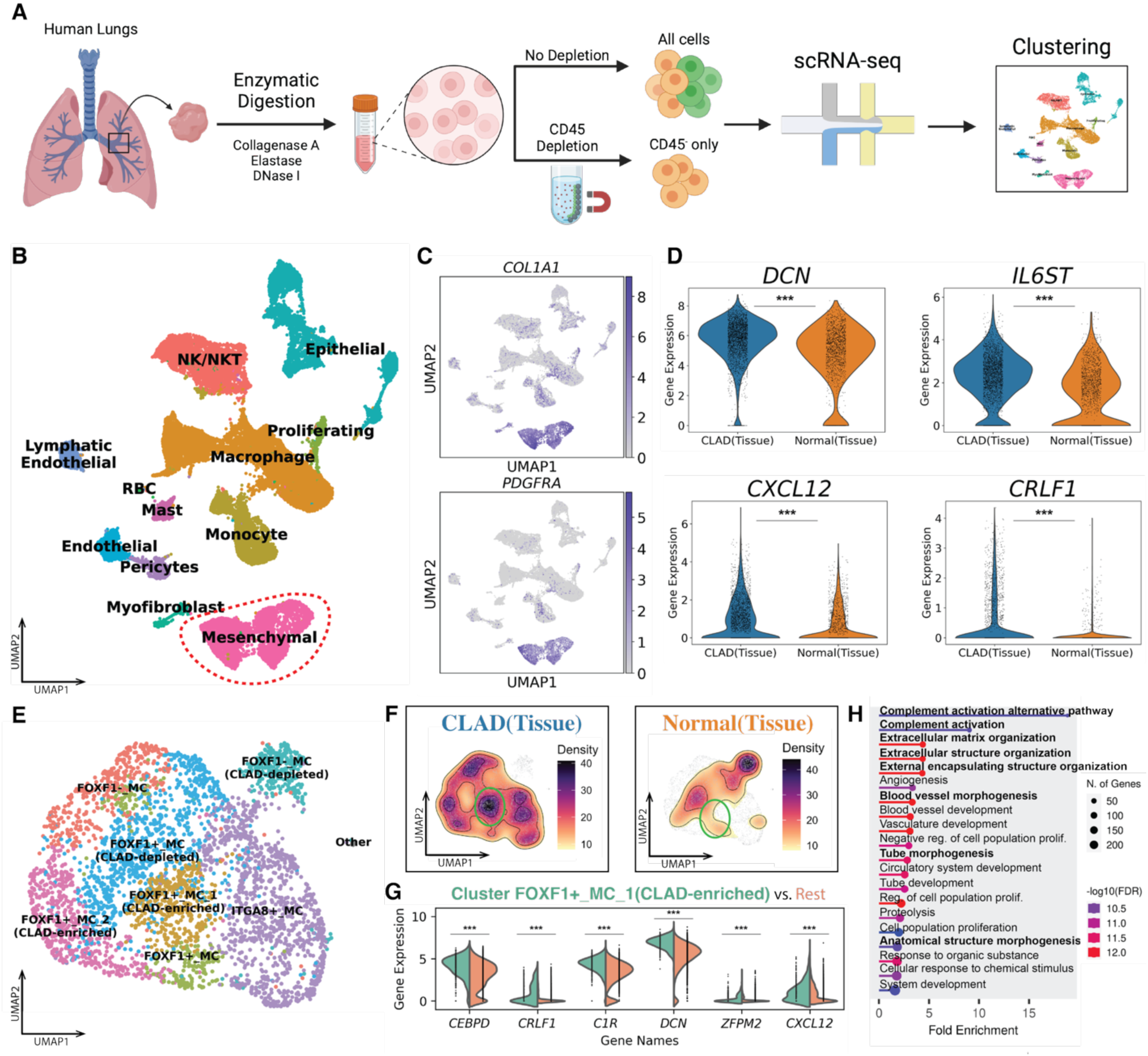
Cultured MCs retain and reflect their *in vivo* phenotype. **(A)** Overview of scRNA-seq sample preparation and analysis. **(B)** UMAP representation of 28,279 cells from both non-depleted and CD45+ depleted samples. Each dot represents a single cell, and cells are colored by cell-types. MCs are circled in a red dashed line. **(C)** UMAP colored by *COL1A1* (upper) and *PDGFRA* (bottom) expression. **(D)** Violin plots depicting the expression levels of *DCN* (upper left), *IL6ST* (upper right), *CXCL12* (bottom left), *CRLF1* (bottom right) across conditions. The width of each plot represents the distribution of expression values. Comparisons are shown between CLAD tissue (blue) and normal tissue (orange). **(E)** UMAP representation of 3,902 tissue MCs. **(F)** Density plots of integrated tissue MCs (left: CLAD tissue; right: Normal tissue). **(G)** Split violin plots showing the expression levels of *CEBPD*, *CRLF1*, *C1R*, *DCN*, *ZFPM2*, *CXCL12*, and *NFIB*. Each violin is split to compare cluster FOXF1+_MC_1 (CLAD-enriched) with the rest of the MCs, with the width representing the distribution of expression values. **(H)** GO enrichment analysis showing upregulation of complement activation-related and ECM-related genes in CLAD tissue MCs. (***p ≤ 0.001)

After quality control filtering, we obtained 28,279 cells from both non-depleted and CD45+ depleted samples. We performed integration and unsupervised clustering using LIGER (30, 31), and identified non-immune cell types including mesenchymal, endothelial, epithelial cells, pericytes, and myofibroblast as well as immune cell types including macrophages, monocytes, mast cells, nature killer (NK) / nature killer T-cells (NKT), red blood cells (RBCs), and proliferating cells (**Figure 7B**). A total of 4,254 tissue MCs were identified based on the expression of *COL1A1* and *PDGFRA* (**Figure 7C**). Differential expression analysis between CLAD tissue MCs and healthy controls revealed 5,263 upregulated genes and 5,130 downregulated genes in CLAD tissue MCs. To assess the transcriptional similarity between CLAD tissue MCs and CLAD cultured MCs, we compared these differentially expressed genes with the previously identified CLAD signature in Figure 2 from the logistic regression model. Notably, 9 out of 17 upregulated genes in CLAD cultured MCs were also significantly upregulated in the CLAD tissue MCs (**Figure 7D; Additional file 1: Figure S7B**). Of these, 5 genes (*DCN*, *IL6ST*, *CXCL12*, *CRLF1*, and *NTNG1*) were predicted as the downstream targets of CEBPD based on the eGRN, and 4 genes (*IL6ST, CXCL12*, *NTNG1*, and *NFIB*) had previously been shown to be downregulated after *CEBPD* silencing. Additionally, 6 out of 30 downregulated genes in CLAD cultured MCs were also significantly downregulated in the CLAD tissue MCs, with 1 gene (*ID1*) previously shown to be downregulated after *CEBPD* silencing (**Additional file 1: Figure S7C**). These results suggest that MCs *in vivo* shared key transcriptomic signatures with their cultured counterparts.

To obtain a more fine-grained resolution of cell states in tissue MCs, we performed sub-clustering (**Figure 7E**). This analysis identified a sub-cluster, FOXF1+_MC_1 (CLAD-enriched), which was specifically enriched in CLAD samples (**Figure 7F; Additional file 1: Figure S8A**). Importantly, this sub-cluster exhibited the highest expression of *CEBPD* compared to other clusters, consistent with the fibrotic state observed *in vitro* (**Figure 7G**). Additionally, 5 genes (*CRLF1*, *C1R*, *DCN*, *ZFPM2*, and *CXCL12*), predicted as downstream targets of CEBPD based on the eGRN, were significantly upregulated in this sub-cluster (**Figure 7G**). Furthermore, 2 CLAD-upregulated signature genes (*NFIB* and *COL3A1)* were also significantly up-regulated (**Additional file 1: Figure S8B**). To further assess the enrichment of our signature genes, we performed Gene Set Enrichment Analysis (GSEA) (41, 42). The analysis revealed significant enrichment in the upregulated signature with a normalized enrichment score (NES) of 1.97.

Although the whole downregulated gene set was not significantly enriched, some of the genes still were significantly downregulated in this sub-cluster (**Additional file 1: Figure S8C**). GO enrichment analysis of the differentially expressed genes in this sub-cluster revealed significant upregulation of biological processes related to extracellular matrix (ECM) organization, ECM structural organization, and external encapsulating structure organization, suggesting their fibrotic phenotype and aligning with the top downregulated GO terms observed after *CEBPD* silencing. Additionally, several GO terms associated with morphogenesis were enriched, consistent with those observed in the fibrotic state of MCs *in vitro*. These results suggest this sub-cluster may represent the *in vivo* counterpart of the fibrotic state identified *in vitro*.

## Discussion

Fibrosis is an important pathogenic finding in chronic allograft failure across all solid organ transplants. However, targeting fibrogenesis continues to be elusive with fibrotic MCs demonstrating multiple dysregulated pathways. In this study, we leveraged genetic multiplexing and single-cell multi-omics to provide an unbiased examination of distinct MC gene expression and chromatin accessibility signatures in lung allografts undergoing early inflammatory and late fibrotic processes. Genetic multiplexing—pooling cells from multiple donors and demultiplexing them based on natural genetic variation—enabled us to reduce batch effects and increase the number of cells and donors profiled in our study. Simultaneous profiling of gene expression and chromatin accessibility within the same cells was essential for linking transcriptional changes with their upstream regulatory elements. Computational approaches integrating these chromatin accessibility and gene expression measurements into an enhancer-driven gene regulatory network were also key to identifying disease-associated molecular changes. Through this integrative approach, we identified unique gene signatures in CLAD patient-derived MCs and uncovered interesting molecular features in these MC subtypes with the enrichment of transcription factors such as CEBPD, SOX4, and FOXP2 in the fibrotic MC state. The gene regulatory networks we identified suggest that these transcription factors may play pivotal roles in modulating gene expression programs critical for fibrogenesis and immune interactions in CLAD.

Our unbiased analysis of the global transcriptome of MCs isolated from transplanted lungs reveals a distinct MC transcriptional signature in patients with CLAD, with CLAD-derived MCs exhibiting a pronounced shift towards a fibrotic state. CLAD-MCs could be differentiated from non-CLAD with 99.2% accuracy and specific upregulated and downregulated genes were identified. An important control was our comparison of MCs isolated from patients early post-transplant, where differentiation between the two groups was not evident. CLAD emerges over time, with the incidence of CLAD increasing over time post-transplant. The early post-transplant group included patients who developed CLAD later in the course of their transplant, with the average time to CLAD of 3.5 years (10). MCs isolated at the early stage could not be classified with high accuracy into diseased and non-diseased groups, and these cells were compositionally similar to each other and the non-CLAD later post-transplant control group. This definitive demonstration of an acquired stable fibrogenic transformation in CLAD MCs offers further novel insight into our understanding of immune-mediated fibrotic processes, where immune regulation after disease onset fails to slow disease progression.

A key finding was the identification of transcription factor CEBPD as a key regulator of the fibrotic state of MCs in CLAD. CEBP-family members, specifically CEBPD, were dominant TFs identified in analyses of chromatin accessibility profiles of CLAD-MCs and their binding motifs using motif scan. Its specificity to the CLAD state was suggested by a parallel analysis of early time post-transplant MCs, where CEBPD was not noted among their enriched motifs. Gene regulatory analysis combining single-cell transcriptomic and epigenomic features nominated CEBPD as a key regulator of CLAD-enriched fibrotic state, with both the most state-specific expression in the fibrotic state and the most predicted targets. Further, *in vivo* significance was shown by a similar profile in MCs directly analyzed from a CLAD lung homogenate with significantly higher expression of these genes. The silencing of *CEBPD* via siRNA in CLAD-MCs provided functional support for its regulatory role. The bulk RNA-seq data demonstrated that silencing *CEBPD* downregulated key fibrotic genes and shifted the overall gene expression profile of CLAD MCs towards a less fibrotic state. Key CLAD-associated genes including *CXCL12* and *IL6ST* were found to be direct downstream targets of CEBPD. While very few studies have investigated CEBPD in MCs in the context of fibrosis, CEBPD has been demonstrated to be the master TF regulating extracellular matrix proteins including fibronectin, and several collagens in glioblastoma (43). CEBPD was shown to be a mediator of hypoxia responses and critical proteins of TGFβ signal pathway, *TGFBI* and a component of integrin receptor *ITGA5* were also proposed to be direct targets of CEBPD (43). Extracellular matrix structural constituent was also the most significant downregulated GO term in our analysis of *CEBPD*-silenced CLAD MCs. Among the downregulated ECM genes were periostin (*POSTN*) and thrombospondin1 (*TSP1*), both key players in fibrosis along with fibrillin 1 and 2 (*FBN1/2*), and multiple collagens. Future studies are needed to investigate the cell-specific role of CEBPD in fibrosis and to validate a requisite role for CEBPD in CLAD pathogenesis using *in vivo* models. However, regulation of a master TF to reverse the fibrogenic state of MCs represents a novel and promising therapeutic target for mitigating fibrosis, and our investigations suggest CEBPD to be one such putative regulator.

Among the genes identified in this unbiased approach are targets that are being increasingly recognized across other fibrotic diseases and have a direct link to prior investigation of the mechanisms of lung allograft fibrosis. *CXCL12, IL6ST,* and *CRLF1* were found to be: (i) upregulated in CLAD MCs in an unbiased genomic analysis; (ii) were confirmed to be downregulated regulated by *CEBPD* silencing by RT-qPCR, and importantly; (iii) shown to be significantly upregulated in fibrotic MCs in single cell examination of CLAD lung tissue. Interestingly, all three of these pathways link to our previously described signaling pathways in CLAD MCs. We have recently characterized CXCL12 expression by *Foxf1*+ MCs and its role as a regulator of graft ASC niche in RAS (44). IL6ST or glycoprotein 130 (gp130), the universal signal-transducing β-receptor subunit for all IL-6-related cytokines, allows for IL-6 trans-signaling in cells lacking IL-6 receptor (45), and we have demonstrated that IL6 trans-signaling regulates both MC fibrotic differentiation (17) and CXCL12 expression (44). Interestingly, others have shown that GP130 upregulation can create an autocrine activation loop involving IL-6/sIL-6R (46). Cytokine receptor-like factor 1 (CRLF1) has been implicated in fibrosis in a variety of tissues (47–49), with specific relevance to our previous findings of increased mTORC2 signaling and upregulation of critical mTORC2 component Sin1 in CLAD MCs. CRLF1 is recognized as a novel mTORC2 component that strengthens the interaction between Sin1 and AKT, enhancing AKT Ser473 phosphorylation (50).

The diagnosis of CLAD is made clinically using spirometric decline, as transbronchial biopsies are not sensitive enough to offer histologic confirmation. CLAD is classified into clinical phenotypes based on spirometric decline pattern and radiographic features (10). While some patients are classified into the two predominant clinical phenotypes of bronchiolitis obliterans syndrome and restrictive allograft syndrome, others fall into indeterminate or mixed category. Fibrosis of the allograft is common to all phenotypes with fibrosis emanating from bronchovascular bundles being an important central feature in majority of the rejecting grafts. Our previous work on MCs in BAL has offered a window into fibrogenesis in a transplanted lung. We have shown that increased numbers of MCs in BAL, as quantified by CFU assay, precede the development of CLAD, with mesenchymal CFUs noted to be a strong predictor of CLAD onset (14). Furthermore, high mesenchymal CFUs in BAL at CLAD onset were associated with poor survival and restrictive allograft syndrome diagnosis (15). The CLAD MCs utilized in this multiomic study were from patients with diverse clinical CLAD phenotypes, and the present study does not have the power to address the specific correlation with CLAD phenotypes. Further work is needed to understand if epigenetic changes in CEBPD mark a specific clinical or molecular phenotype and will require a large study sample with outcome analysis. Our study included MCs from lungs after CLAD onset or very early post-transplant. While this allowed us to determine if CLAD-associated changes are acquired over time, a longitudinal study of MCs at various time points in patients with variable post-transplant complications, like acute rejection or infection episodes, can shed more light on the time course of emergence of this fibrogenic phenotype and the preceding insults.

In conclusion, our single-cell multi-omics analysis revealed key molecular signatures and regulatory networks associated with fibrosis and CLAD. The identification of a distinct transcriptomic signature and a skewing towards a more homogeneous fibrotic state in CLAD-MCs underscores the acquisition of a stable fibrogenic transformation in graft MCs that drives CLAD. Our work is the first to identify transcription factor CEBPD as a master regulator of fibrotic cell state and links it to key downstream genes in MCs. Future studies should focus on further dissecting the regulatory mechanisms at play and exploring combinatorial therapeutic strategies to effectively combat fibrosis and chronic rejection in lung transplantation.

## Methods

### Sex as a biological variable

MCs were cultured from bronchoalveolar lavage fluid samples derived from lung transplant patients of both female and male sex (Supplemental Table 1). None of the experiments nor analyses were limited to samples from either sex. As such, sex was not evaluated as a biological variable in the experiments.

### MC isolation from human lung allografts and cell culture

BAL samples were obtained from human lung transplant recipients under a protocol for human studies approved by the University of Michigan Institutional Review Board. MCs were isolated from BAL as previously described (6, 7, 11, 12, 14, 16–18). Briefly, all cells isolated from BAL fluid were pelleted by centrifugation (400 x g for 5 minutes) and were maintained in adherent culture conditions. MCs were identified by their growth as colony-forming units (CFUs) with fibroblastoid morphology. Cells were maintained in culture in high-glucose 1X DMEM (Gibco, Thermo Fisher Scientific) supplemented with 10% FBS (Avantor), 100 U/ml penicillin/streptomycin (Gibco), and 0.5% Amphotericin B (Gibco), at 37°C in humidified incubators with 5% CO2 for no less than one week and no more than three weeks following initial sample processing before being passaged by trypsinization. Individual bronchoalveolar lavage samples and subsequent MCs isolated were treated as separate cell lines. Cells from passages three through six were used for all experiments. CLAD was defined by a >3-week persistent decline in FEV1 or FVC utilizing ISHLT guidelines and as previously described (10, 51, 52). CLAD phenotyping was performed using the latest ISHLT guidelines (10). At the early time point (<6 weeks post-transplant), samples were collected from patients who had not yet developed CLAD but were later clinically classified as having developed CLAD (n=4) or remained CLAD-free (n=3). At the late time point (>1year post-transplant), samples were obtained from patients with CLAD (n=3) and without CLAD (n=2) (**Table S1**).

### Cell Preparation for single-cell multiome sequencing

MCs were plated on 10cm dishes and grown to full confluence. For 10X Multiome RNA+ATAC-sequencing, nuclear isolation was performed according to demonstrated protocols from 10X Genomics (CG000366, Revision C). Briefly, culture media was aspirated, and plates were washed twice with cold PBS. Cells were isolated using a cell scraper and 5mL of ice-cold PBS into a 15mL conical tube and spun at 500 x g at 4C. The supernatant was removed and the cells were resuspended using 1X lysis buffer containing 10mM Tris-HCl (Invitrogen), 10mM NaCl (Fisher Scientific), 3mM MgCl2 (Promega), 0.1% Tween-20 (Fisher Scientific), 0.1% Nonidet P40 Substitute (Sigma-Aldrich), 0.01% Digitonin (Thermo Fisher Scientific), 1% BSA (Sigma-Aldrich), 1mM DTT (Sigma-Aldrich), and 1U/uL RNasin Plus Ribonuclease Inhibitor (Promega) in Nuclease-Free Water (Invitrogen) and incubated on ice for 5 minutes. Cells were monitored until less than 10% of cells remained unlysed. The pellet was then washed two times with centrifugation at 500 x g for 5 minutes at 4C, each time removing the supernatant. Finally, 10,000 target nuclei were resuspended in a diluted nuclei resuspension buffer (10X Genomics), pooling 2-3 cell lines for each submission.

### Single-cell multiome library generation

Single-cell multiome libraries were prepared using the Chromium Next GEM Single Cell Multiome ATAC + Gene Expression kit (10X Genomics) according to manufacturer’s instructions. ATAC sequencing, library preparation, and 3’ gene expression was performed with 50,000 reads per cell depth. Sequencing was performed by the Advanced Genomics Core at the University of Michigan using a NovaSeq 6000 (Illumina) platform.

### Single-cell multi-omic data preprocessing and quality control

Cell Ranger ARC pipeline (version 2.0.0) with default parameter settings was used to process the raw Chromium Single Cell Multiome ATAC + Gene Expression sequencing data. During alignment steps, reads were mapped to the human reference genome (GRCh38) provided by 10x Genomics. The filtered count matrices output by the pipeline were used for further analysis. Quality control steps were applied separately to the RNA and ATAC modalities, each using specific quality control metrics. For the RNA modality, low-quality cells with fewer than 200 unique genes or more than 20% mitochondrial reads were removed. For the ATAC modality, we assess the data quality using four statistics: Fraction of Reads In Peaks (FRIP), Transcription Start Site (TSS) enrichment score, log number of unique fragments, and duplication rate (**Additional file 1: Figure S4**). Low-quality cells with fewer than 0.3 FRIP, 4 TSS enrichment score, or 3.3 log number of unique fragments were removed.

### Genotyping and demultiplexing of single-cell multiome sequencing data

Cultured MCs from each cell line submitted for multiome sequencing were grown to confluence before their whole cell DNA was isolated using the DNeasy Blood and Tissue Kits (Qiagen) according to manufacturer’s instructions. DNA concentrations were measured via absorbance at 260/280nm using a NanoDrop 2000 (ThermoFisher) prior to submission to the Advanced Genomics Core (AGC) at the University of Michigan. Demultiplexing of all 16 samples and associated pooled nuclei was conducted using an INF Multi-Ethnic Global-8 Kit (Illumina) genotyping kit. A set of SNPs was first obtained, then allelic expression of these variants in each cell was extracted using the read pileup approach (dsc-pileup from popscle for demuxlet or cellsnp-lite mode 1a for vireo). scRNA-seq and scATAC-seq were demultiplexed using demuxlet and Vireo separately. We observed doublet overestimation by both demultiplexing softwares (demuxlet and vireo; **Additional file 1: Figure S5**) in the RNA modality which was likely due to ambient RNA contamination. Therefore, the final assignment of cell identity was determined by taking into consideration both assignments from the RNA and ATAC modalities. Cells that have the same cell assignment between two modalities were kept. In addition, if a cell was identified as doublet in RNA, but not in ATAC, we used the cell assignment from its ATAC modality. We excluded cells that were ambiguous or identified as doublets in both modalities.

### Logistic regression classifier training

The scRNA-seq gene expression matrix was utilized and trained logistic regression classifiers for CLAD vs. non-CLAD. We used logistic regression with an elastic-net penalty, a popular approach for extracting a small set of features that can generalize well despite a relatively small number of samples. Elastic net regression uses a linear combination of the L1 penalty (lasso) and the L2 penalty (ridge), leading to sparse selection of a subset of features without arbitrarily choosing among correlated predictors (53–55). Scikit-learn (version 1.2.2) was used to perform the preprocessing and training. To build our signature from genes with appreciable expression levels, we filtered for genes with > 5000 unique molecular identifiers (UMIs) across all cells. Additionally, to prioritize genes with specific expression in mesenchymal cells, we filtered out genes with a total number of UMI counts exceeding 30,000 in non-mesenchymal cell types. Genes encoded on the mitochondrial chromosome, ribosomal protein genes, and genes encoded on sex chromosomes were additionally removed. A total of 5022 genes for CLAD vs. Non-CLAD genes remained after filtering and were used to train the logistic regression model. Next, counts were normalized by the total expression per cell using the *normalize* function. The normalized counts were then log-transformed. Given single-cell sequencing generated different numbers of cells per condition resulting in a skewed class distribution, upsampling was applied to create balanced class distributions. The minority class was oversampled to match the number of cells in the majority class using the RandomOverSampler function from the imbalanced-learn package (version 0.10.1). The preprocessed data were partitioned into training and testing sets in a ratio of 9:1. Testing data were used to evaluate the model performance and were held out from training. Before training the model, each gene feature in the training dataset was normalized to zero mean and unit variance using the StandardScaler function. A logistic regression model with Elastic-Net penalty was then trained using the SGDClassifier (loss=’log’, penalty=’elasticnet’, l1_ratio=0.5). Five-fold cross-validation was performed to select the alpha and evaluate the model performance using the GridSearchCV function. To further validate gene signatures from our classifier, we performed real-time quantitative PCR (RT-qPCR) on a selected set of genes.

### mRNA Isolation, cDNA Synthesis, and Gene Expression Analysis

Following siRNA-mediated transfection, cells were washed two times with PBS (Gibco) and lysed using RLT-buffer (Qiagen) supplemented with 2-Mercaptoethanol (Fisher Scientific). Total mRNA was isolated immediately using RNeasy Plus Kit (Qiagen) according to the manufacturer’s instructions. RNA quantity was assessed using a NanoDrop 2000 (Thermo Fisher). cDNA synthesis was performed using a High-Capacity cDNA Reverse Transcriptase Kit (Applied Biosystems) according to manufacturer’s instructions. cDNA synthesis was conducted on a 96-well Veriti Thermocycler (Applied Biosystems). After cDNA synthesis, gene expression analysis with TaqMan gene expression master mix (Applied Biosystems) was performed with TaqMan assays (Applied Biosystems) for *ACTB* (Hs01060665_g1), *CXCL12* (Hs00171022_m1), *IL6ST* (Hs00174360_m1), *CRLF1* (Hs00191064_m1), *DCN* (Hs01081921_m1), *CEBPD* (Hs00270931_m1). All RT-qPCR assays were run on a QuantStudio3 Real Time PCR machine (Applied Biosystems).

### scATAC-seq data analysis

ArchR (version 1.0.1) was used for the inter-sample analysis of scATAC-seq data. Pseudo-bulk replicates were first generated to obtain measurements of statistical significance using the addGroupCoverages function with default parameters.

Reproducible merged peak sets were generated using the addReproduciblePeakSet function, which first called peaks per dataset using MACS2 and then iteratively merged overlapping peaks across datasets. Marker peaks were identified using the getMarkerFeatures function and annotated using the peakAnnoEnrichment function to obtain enriched motifs. The pairwise Wilcoxon testing was performed between CLAD and non-CLAD groups and the cutoffs were false discovery rate ≤ 0.05 and absolute log2 fold change ≥ 1.

### scRNA-seq data integration

scRNA-seq data matrices after quality control were integrated using PyLiger (version 0.0.1)(31), a Python version of our previously developed algorithm, linked inference of genomic experimental relationships (LIGER) (30). LIGER is a widely used package for single-cell multi-omic data integration. Briefly, it employs integrative non-negative matrix factorization (iNMF) to identify shared and dataset-specific metagenes of cellular variation. Before integration, scRNA-seq data were first analyzed with standard preprocessing steps required by LIGER including normalization, highly variable gene selection, and scaling to unit variance. Joint matrix factorization was performed on the normalized and scaled datasets followed by quantile normalization. We then clustered the cells using the Leiden algorithm. Given we noticed biological variation among cells from different donors, we performed LIGER by donor; the count matrices were split into donor-specific count matrices based on donor identity assigned during genetic demultiplexing. The density plots were visualized using customized R code. The compositional changes were evaluated by calculating the proportion of cell types within each sample and visualized using the *boxplot* function from scCODA(56).

### Gene ontology enrichment analysis

The Wilcoxon rank-sum test was conducted using the *runWilcoxon* from LIGER to identify differentially expressed genes (DEGs). We filtered DEGs based on an adjusted p-value < 0.05 and an absolute log fold-change > 2 to identify both up-regulated and down-regulated genes. Gene Ontology (GO) enrichment analysis and network visualization were subsequently performed on the DEGs using ShinyGO (version 0.77) (57). Default parameter settings in ShinyGO were applied, including an FDR cutoff of 0.05, a minimum pathway size of 2, a maximum pathway size of 5000, and a display of the top 20 enriched pathways.

### RNA velocity analysis

Spliced and unspliced counts of each sample were generated using Velocyto (58). To correct the potential batch and donor differences and prepare input for VeloVAE, both RNA and ATAC matrices as well as spliced and unspliced counts were integrated using Seurat’s *FindIntegrationAnchors* and *IntegrateData* functions. The corrected matrices were used to train a VeloVAE model (35). The velocity stream plot was visualized using the *velocity_embedding_stream* function from scVelo.

### Enhancer-driven gene regulatory network analysis

SCENIC+ was employed to build an enhancer-driven gene regulatory network (36). The SCENIC+ workflow consists of three analysis steps: (i) using the pycisTopic module to identify candidate enhancers; (ii) using the pycisTarget module to identify enriched TF-binding motifs; and (iii) using the SCENIC+ module to link TFs to candidate enhancers and target genes.

*pycisTopic.* The consensus peak set generated by ArchR was used to identify candidate enhancers. Topic modeling was performed using the *run_cgs_models* function from pycisTopic, with the number of topics tested set to 5, 10, and 15.

*Motif enrichment analysis.* In the second step, the *run_pycistarget* function from pycisTarget was used to perform motif enrichment analysis with default parameters. The precomputed cisTarget database was downloaded from the pycisTarget website.

*Inferring eGRNs using SCENIC+.* Finally, the run_scenicplus function from the SCENIC+ module was used to construct eGRNs. We used default parameters. The heat map/dotplot was generated using the heatmap_dotplot function on significant eRegulons. TF cooperation activities were calculated using the jaccard_heatmap function. The network was visualized using the *plot_networkx* function.

### Western blot

*Protein isolation and immunoblotting.* MCs were plated and grown to 50% confluence in 60-mm dishes prior to serum starvation for 24 h. Total protein was collected from treated or untreated cells by whole cell lysate, lysed using Cell Lysis Buffer (10X, Cell Signaling) diluted to 1X with water and supplemented with 0.1M phenylmethylsulfonyl fluoride (PMSF) in ethanol (Sigma-Aldrich) and EDTA-free Halt Phosphatase Inhibitor Cocktail (Thermo Fisher Scientific). Whole cell lysates were then centrifuged (17,000 x g for 10 minutes) at 4°C. Lysate concentration was determined using a Pierce Coomassie Plus Assay Kit (Thermo Fisher Scientific) and analyzed via spectrophotometry (Thermo Scientific, BioMate 3 Spectrophotometer). Lysates were separated on 4–12% gradient Bis-Tris gels (Invitrogen) prior to immunoblot analysis for CEBPD (Cell Signaling), GAPDH (Santa-Cruz). HRP-conjugated anti-mouse and anti-rabbit secondary antibodies were A8924 (Sigma-Aldrich) and A0545 (SigmaAldrich), respectively. Immunoblots were imaged using a ChemiDoc MP (Bio-Rad). Densitometry analysis completed using ImageJ (NIH, Bethesda, MD).

### CEBPD knockdown

*siRNA-mediated silencing transfection.* We transiently transfected CLAD MCs at 60% confluence with 100nM of ON-TARGETplus SMARTpool siRNA targeting either: *CEBPD* (Horizon) or a non-targeting control (Horizon). Briefly, cells were plated to 60% confluence in 60mm dishes before transient transfection in reduced serum containing media Opti-MEM (Gibco) with respective pools of siRNAs utilizing Oligofectamine (Invitrogen) for 24hrs. Following transfection, cells were maintained in serum-free medium for either 24 or 48 hrs prior to mRNA isolation or protein harvesting, respectively.

### Bulk RNA-seq preparation and sequencing

CLAD MCs treated with either scrambled (non-targeting) control siRNA or CEBPD siRNA were harvested 24h post-transfection. Total RNA was isolated using the RNeasy Mini Plus Kit (Qiagen) according to the manufacturer’s instructions. RNA was quantified via measuring absorbance at 260/280nm using a NanoDrop 2000 (ThermoFisher) prior to submission to the Emory Integrated Genomics Core (EIGC). After RNA quality assessment was performed using a TapeStation system (Agilent Technologies), all mRNA was sent to a sequencing company (Novogene, China). Bulk RNA-seq (Eukaryotic RNA-Seq, non-directional, stranded library) was conducted by Novogene using the Illumina NovaSeq PE150 platform that employs a paired-end sequencing technology yielding 150-bp read lengths, with all raw reads first cleared of adapter sequences.

### Bulk RNA-seq data alignment and differential expression analysis

Raw RNA-seq reads from both treated and control groups were aligned to the GRCh38 reference genome using STAR (v.2.7.11a) with default parameters. STAR produced sorted BAM files, which were used as the input for HTSeq-count (v.2.0.8) to generate gene-level read counts. The raw gene counts were then imported into DESeq2 (v.1.40.2) for differential expression analysis. DESeq2 was used to measure the effect of the condition (scrambled siRNA vs. CEBPD siRNA), controlling for donor differences.

We used DESeq2 to normalize the data to account for differences in sequencing depth and other technical variations, using the median of ratios method. Statistical analysis was performed using the Wald test, and differentially expressed genes were identified using an adjusted p-value threshold of < 0.05. Note that we didn’t set a cutoff for log2FC to keep genes with small but statistically significant expression changes, ensuring that statistically significant but subtle alterations in gene expression were not excluded from downstream analyses. Gene Ontology (GO) enrichment analysis and network visualization were subsequently performed on the DEGs using ShinyGO (version 0.80). Default parameter settings in ShinyGO were applied, including an FDR cutoff of 0.05, a minimum pathway size of 2, a maximum pathway size of 5,000, and a display of the top 20 enriched pathways.

### CLAD and normal human lung samples

Normal human lungs were procured via of Gift of Life (Michigan). CLAD explanted lungs were procured at the time of re-transplantation under an institutional review board-approved protocol. Representative tissue sections from small airway regions were excised using dissection scissors, manually minced, then placed into a digestion solution containing 1mg/mL Collagenase A (Sigma-Aldrich), 1U/mg/mL Elastase (Worthington), and 0.1mg/mL DNase I (Thermo Fisher) in serum-free 1X high-glucose DMEM. Solutions were incubated for 45min-1hr at 37C in a dry incubator on an end-to-end rotator, mechanically disrupted, and incubated a second time for another 15-30min. Cell suspensions were then filtered through 100um strainers and pelleted by centrifugation at 400 x g for 10min. After aspiration of digestion solution and resuspension, cells were counted and either frozen or taken for CD45+ depletion.

### D45^+^ depletion

For CD45^+^ depletion, 8-10 million frozen cells per tube were segregated from lung digest solutions and placed on ice before being resuspended at a concentration of 2×106 cells/0.5mL in 1X serum-free DMEM (Gibco). Cells were then incubated with 1uL/2×106 cells of CD45-Biotin (Thermo Fisher) antibody at 4C for 30min on a rotator. Cells were then pelleted, washed twice, then incubated with High Capacity Magne Streptavidin Beads (Promega) for another 30min at 4C. Following magnetic capture, remaining CD45-cells were counted, then frozen.

### Primary human lung digest cell submission for scRNA-seq

After initial thawing of frozen cells, they were spun down and resuspended in PBS and kept on ice. A small aliquot of cells was taken at time of submission to monitor for cell death. Samples with >70% viability were submitted for 3’ single-cell RNA sequencing as described above, without additional multiomic sequencing steps.

### scRNA-seq data integration of human lung samples

The Cell Ranger ARC pipeline (verison 6.0.1 for non-delpelted samples and version 6.1.2 for depleted samples) with default parameter settings was used to process the raw Chromium Single Cell Gene Expression sequencing data. During alignment steps, reads were mapped to the human reference genome (GRCh38) provided by 10x Genomics. The filtered count matrices output by the pipeline were used for further analysis. Low-quality cells with fewer than 200 unique genes were first removed. scRNA-seq data matrices after quality control were integrated using PyLiger (version 0.0.1) (31). Before integration, scRNA-seq data were first processed with normalization, highly variable gene selection, and scaling to unit variance. Joint matrix factorization was performed on the normalized and scaled datasets followed by quantile normalization. We then clustered the cells using the Leiden algorithm. DEGs are identified using the Wilcoxon rank-sum test via the *runWilcoxon* function from LIGER (adjusted p-value threshold of < 0.05). Note that we didn’t set a cutoff for log2FC.

### Sub-clustering of tissue MC populations

MC populations identified from integration analysis were subset and re-clustering using LIGER (version 2.1.0). Factor loading of each sample was aligned using *centroidAlign* function. DEGs were calculated in the same way as in the scRNA-seq integration.

*Gene ontology enrichment analysis.* DEGs with adjusted p-value < 0.05 and a log fold-change > 1 was used for GO enrichment analysis and gene set enrichment analysis (GSEA). GO enrichment analysis was performed using ShinyGO (version 0.81). Default parameter settings in ShinyGO were applied. GSEA was performed using GSEAPY (version 1.1.3) with default parameters.

## Author contributions

V.L. and J.D.W. conceived the study. P.M. and N.W. performed all experiments. L.L. analyzed single-cell and bulk RNA-seq data. P.M. analyzed RT-qPCR data. L.L., J.D.W., P.M., R.V., and

V.L. wrote the manuscript. All authors read and approved the final manuscript.

## Data Availability

The raw and processed 10X multiome, bulk RNA-seq and 10X scRNA-seq data are available through the Gene Expression Omnibus (GSE281481, GSE281482, and GSE282248).

## Acknowledgments

We thank Yichen Gu and Chen Li for helpful discussions about RNA velocity and 10X multiome analysis.

## Supplemental Information

**Additional file 1.** Supplementary **Figure S1.** Single-cell multiome quality control metrics. **Figure S2.** Performance of logistic regression models on cells from held-out donors. **Figure S3.** Classifying early time point MCs as developed CLAD vs. not developed CLAD from single-cell gene expression. **Figure S4.** Barcode-level scATAC-seq quality statistics. **Figure S5.** Cell assignment from vireo for the early time point MCs, developed CLAD and not developed CLAD. **Figure S6.** Differentially expressed genes in *CEBPD* knockdown experinmet compared to eGRN predicted targets and CLAD gene signatures. **Figure S7.** scRNA-seq analysis between CLAD lung autopsy samples and healthy controls. **Figure S8.** scRNA-seq analysis of tissue MCs.

**Additional file 2.** Supplementary **Tables S1** Patient demographics and transplant statistics.

---

**Figure S1.**
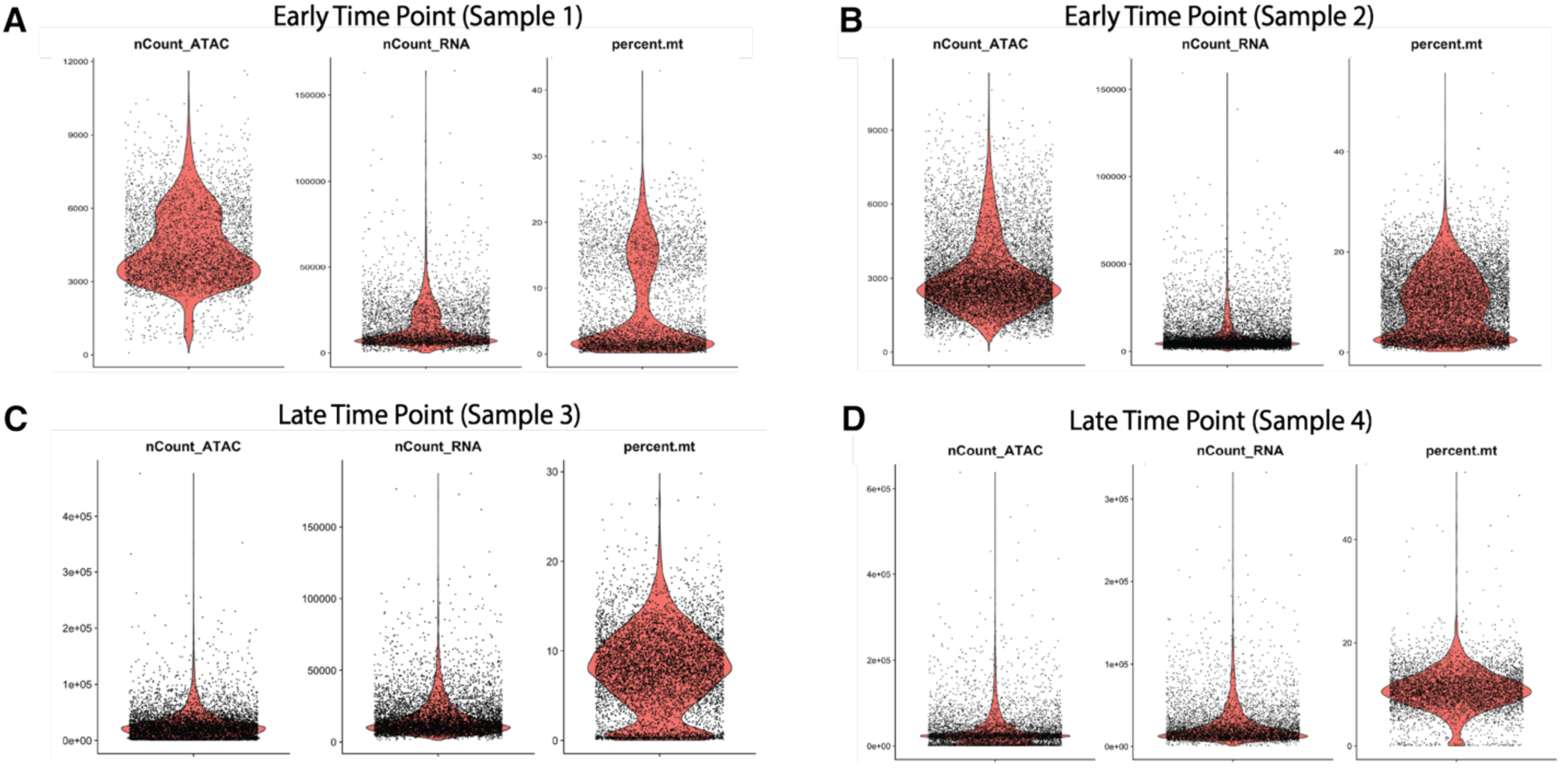
Single-cell multiome quality control metrics.

**Figure S2.**
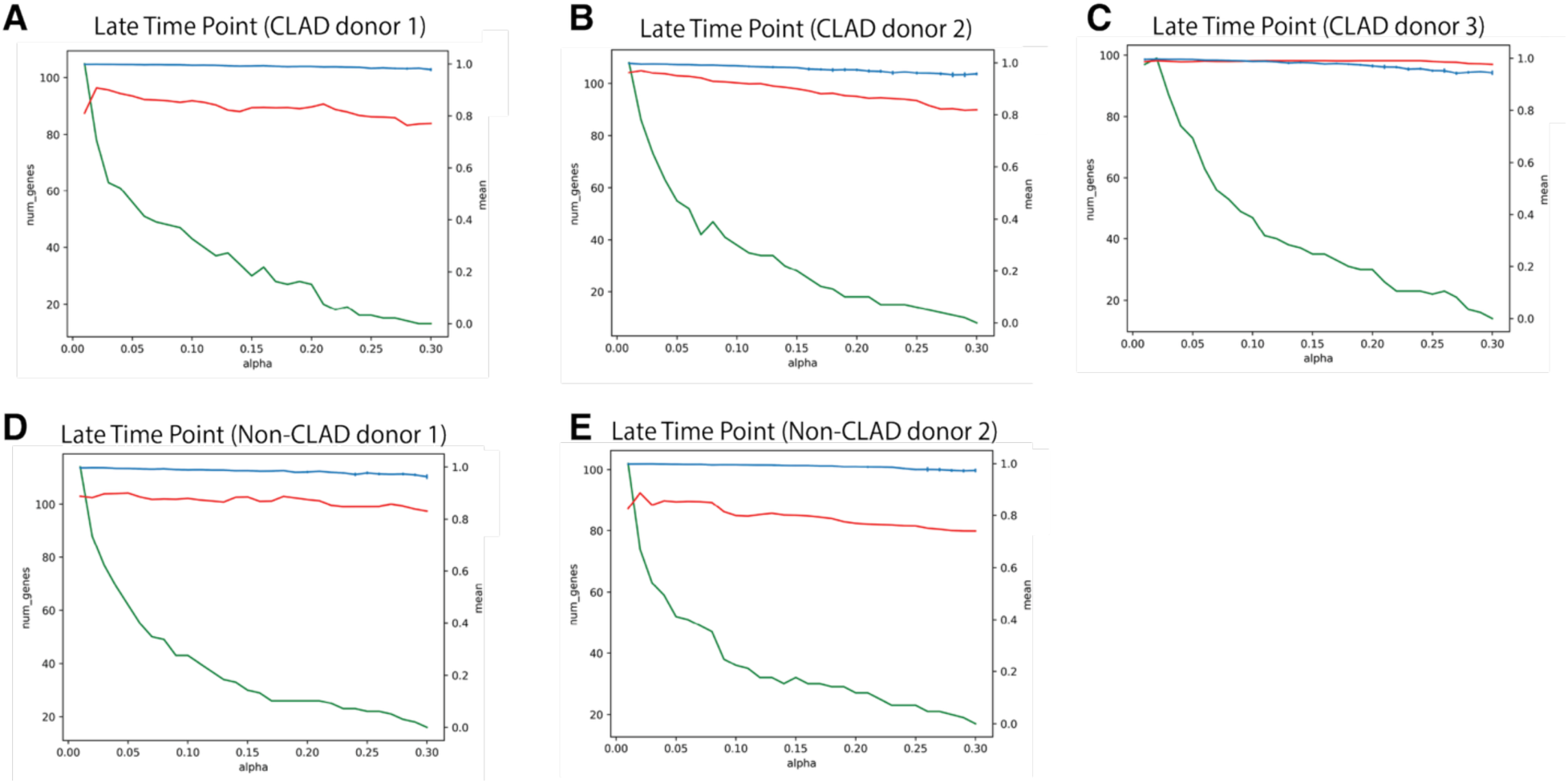
Performance of logistic regression models on cells from held-out donors. The blue line represents the average accuracy of the logistic regression classifiers, calculated using 5-fold cross-validation, with error bars indicating the variance. The green line shows the number of genes, determined by counting the non-zero coefficients from the logistic regression classifiers trained on the test sets. The red line represents the accuracy of the logistic regression classifiers when predicting cells from the held-out donor.

**Figure S3.**
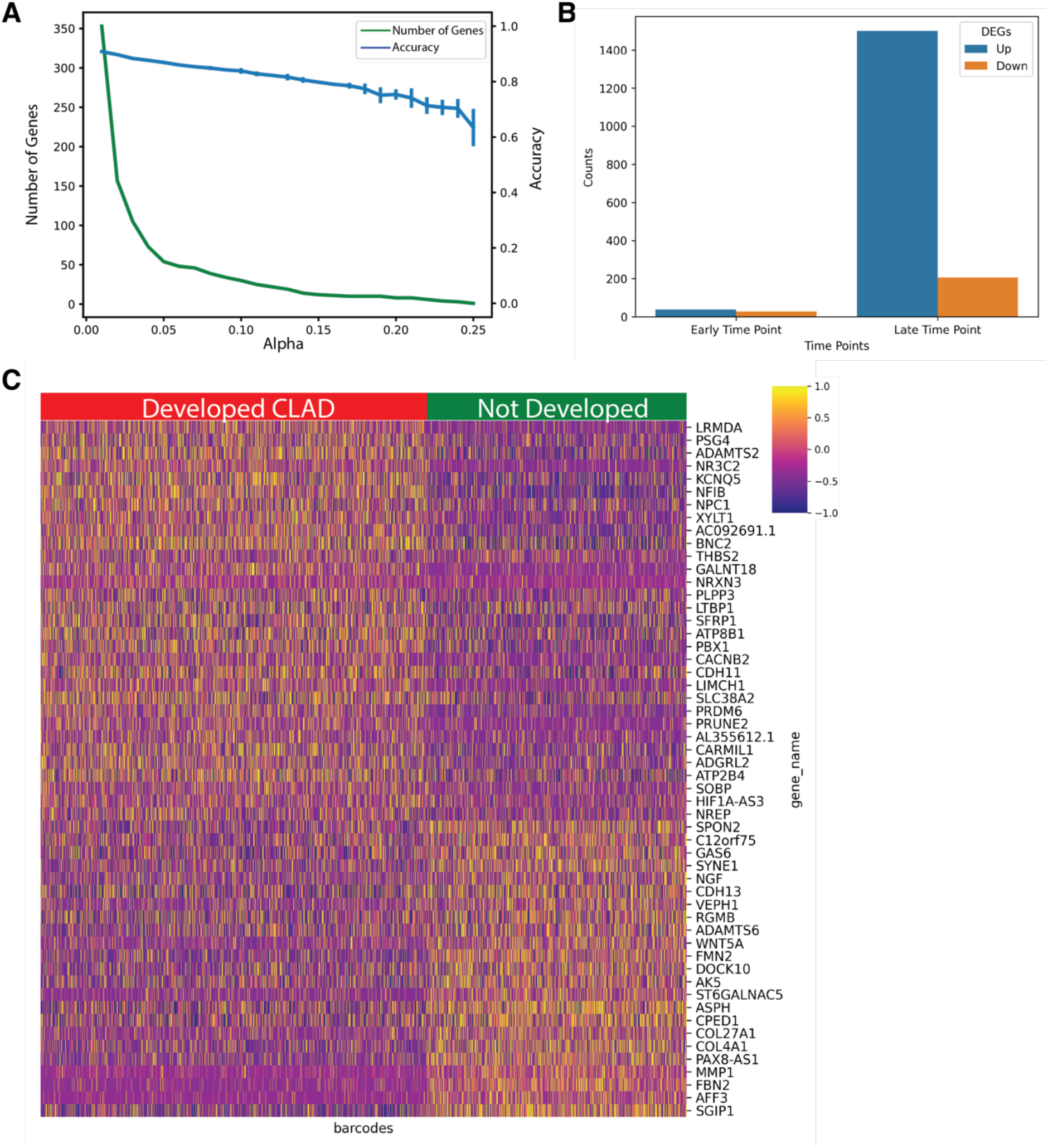
Classifying early time point MCs as developed CLAD vs. not developed CLAD from single-cell gene expression. **(A)** The blue lines indicate average accuracy score of the logistic regression classifiers from 5-fold cross-validation. Error bars indicate the variance in accuracy across folds. The green lines indicate the number of genes with non-zero coefficients from the logistic regression classifiers trained on the test sets. **(B)** The number of differentially expressed genes in early-time-point and late-time-point MCs. Blue bars indicate the upregulated. **(C)** Heatmaps depicting the expression levels of gene signatures in the early time-point MCs constructed from the logistic regression classifiers with α of 0.05. Each column represents a cell, and each row represents one gene colored by gene expression level.

**Figure S4.**
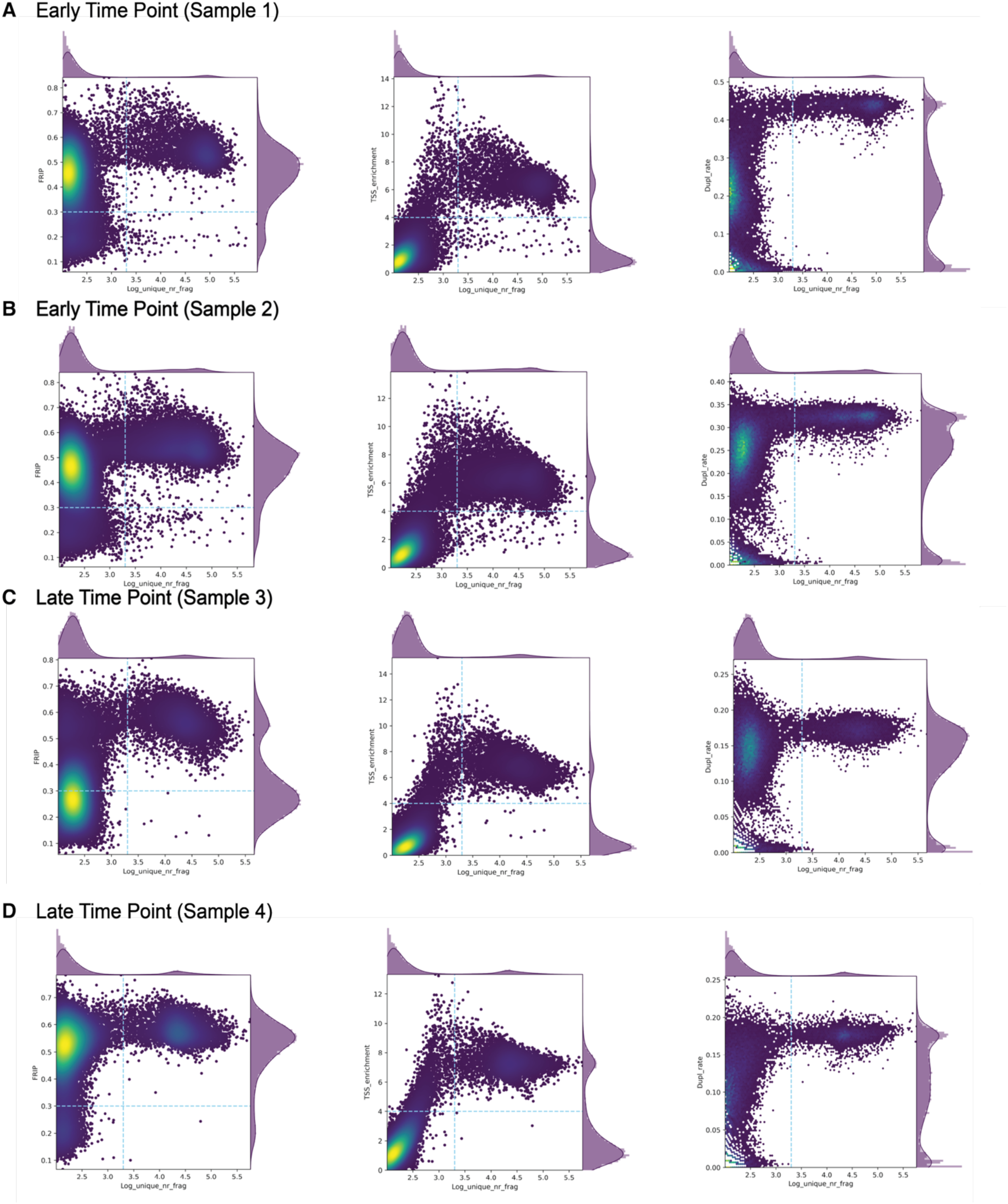
Barcode-level scATAC-seq quality statistics. The x-axis is the log number of unique fragments. The y-axis in the first plot of each sample is Fraction of Reads In Peaks (FRIP). The y-axis in the second plot of each sample is Transcription Start Site (TSS) enrichment score. The y-axis in the third plot of each sample is duplication rate.

**Figure S5.**
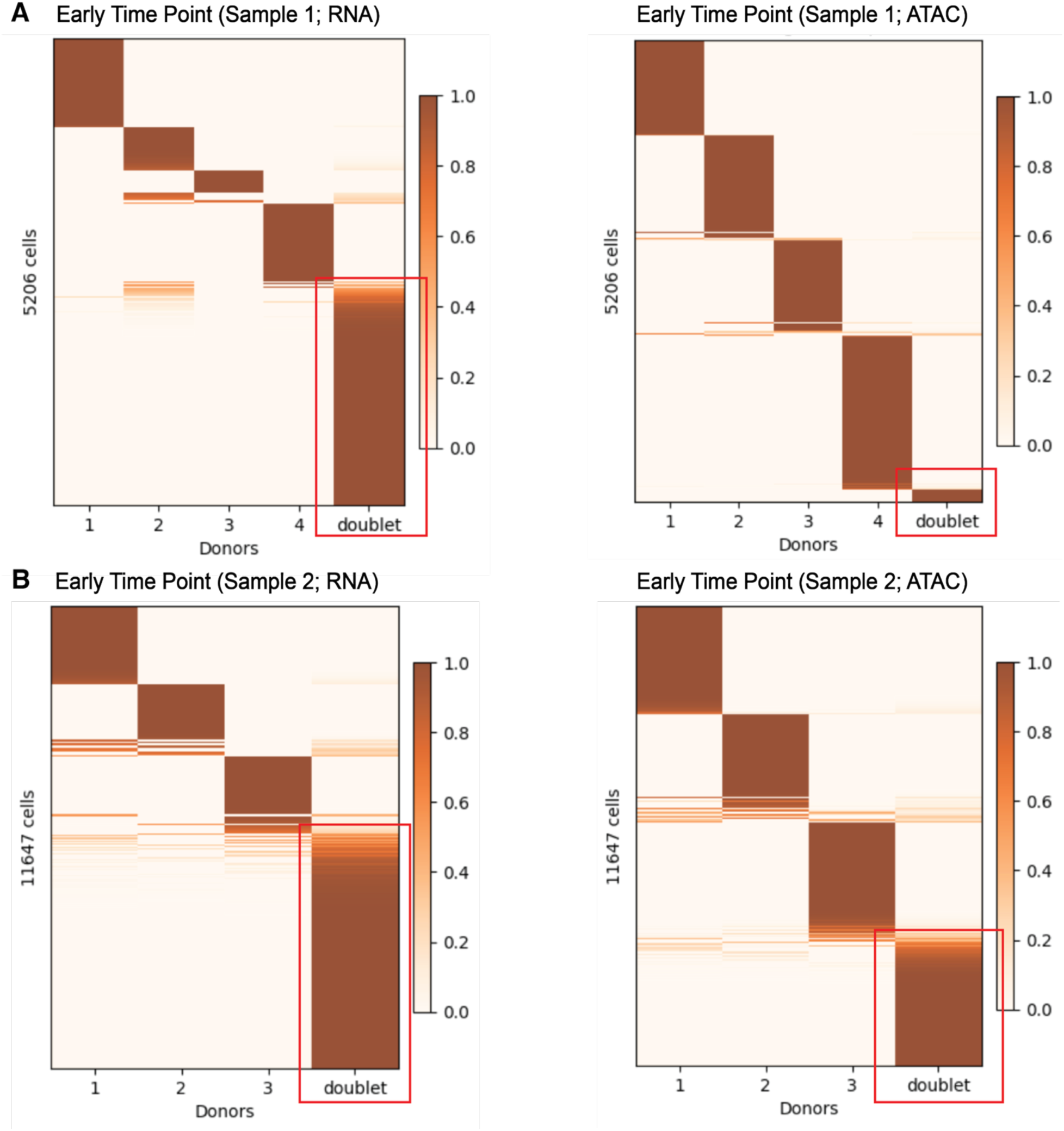
Cell assignment from vireo for the early time point MCs, developed CLAD (A) and not developed CLAD (B). The left plot of each is from the RNA modality and the right plot is from the ATAC modality. The x-axis is the donor id and each line in the y-axis is a cell colored by the probability of assignment to the donor ID. Cells predicted to be doublets by either RNA or ATAC genotype are indicated with a red box.

**Figure S6.**
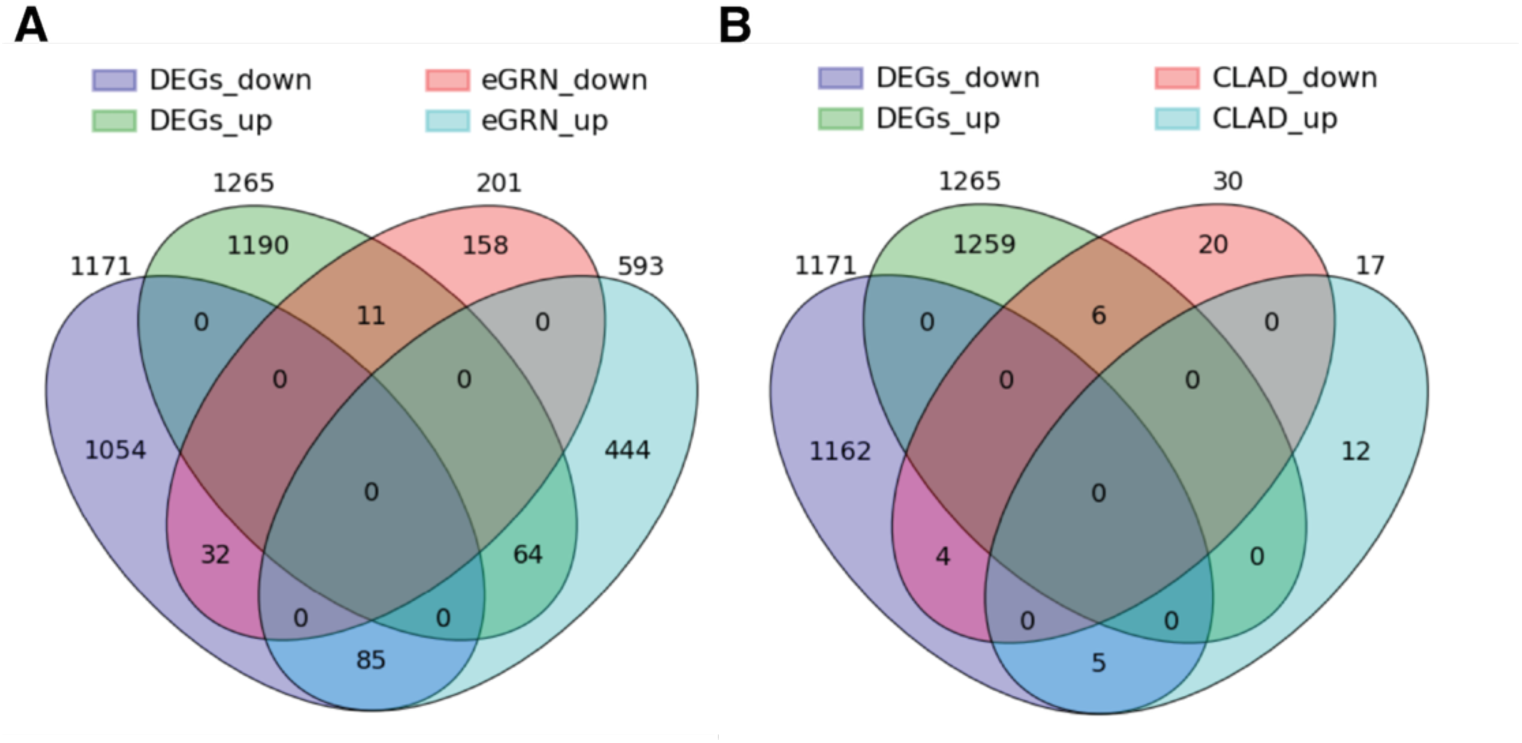
Differentially expressed genes in CEBPD knockdown experinmet compared to eGRN predicted targets and CLAD gene signatures. **(A)** Venn diagram showing the overlap between differentially expressed genes following *CEBPD* silencing and predicted CEBPD targets from the eGRN. **(B)** Venn diagram showing the overlap between differentially expressed genes following *CEBPD* silencing and CLAD gene signatures from Figure 2E.

**Figure S7.**
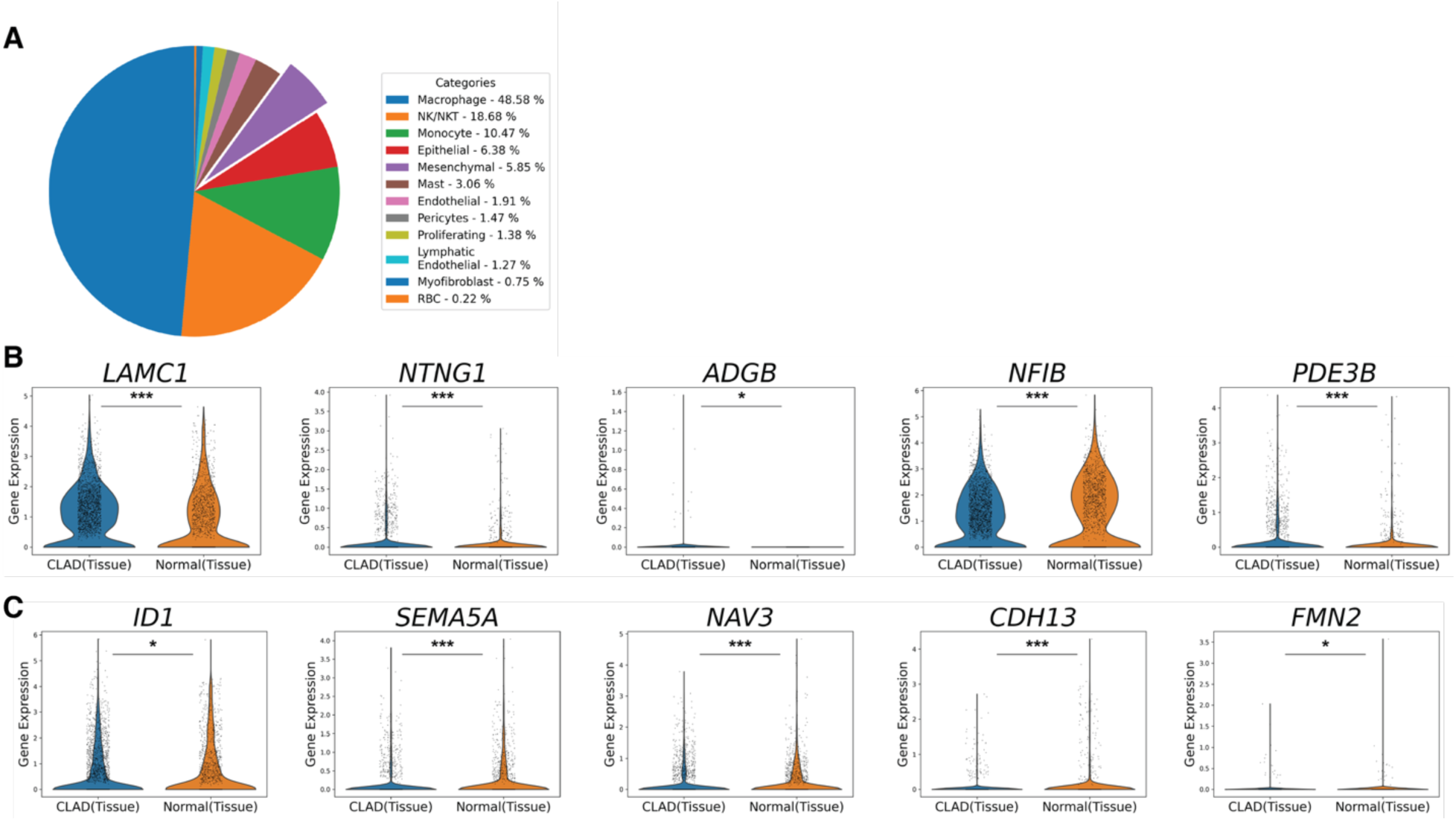
scRNA-seq analysis between CLAD lung autopsy samples and healthy controls. **(A)** Pie chart showing the proportional distribution of each cell-type within non-depleted human lung autopsy samples. **(B)** Violin plots depicting the expression levels of *LAMC1*, *NTNG1*, *ADGB*, *NFIB*, *and PDE3B* across conditions. **(C)** Violin plots depicting the expression levels of *ID1*, *SEMA5A*, *NAV3*, *CDH13*, *and FMN2* across conditions. (*p<0.05; ***p ≤ 0.001)

**Figure S8.**
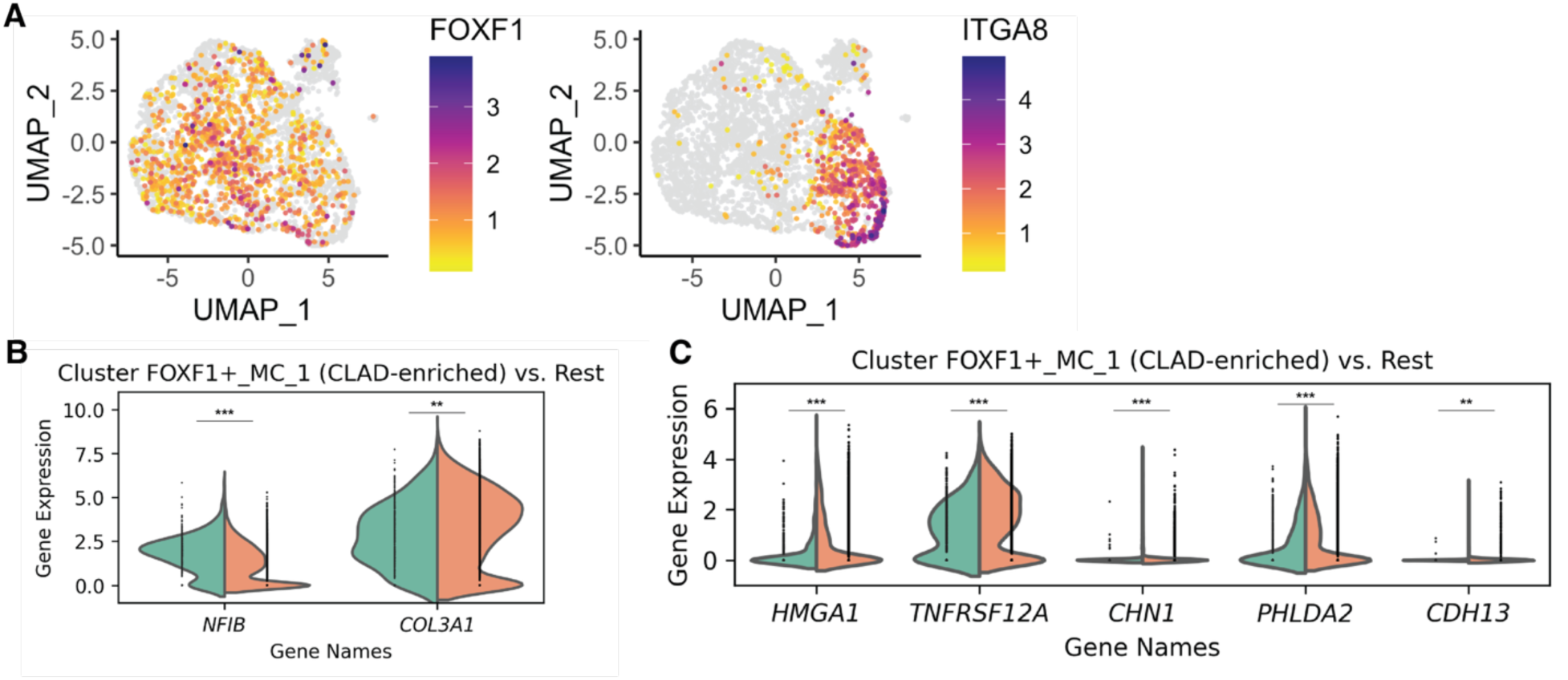
scRNA-seq analysis of tissue MCs. (A,B) Split violin plots showing the expression levels of *NFIB* and *COL3A1*. Each violin is split to compare cluster FOXF1+_MC_1 (CLAD-enriched) with the rest of the MCs, with the width representing the distribution of expression values. **(C)** Split violin plots showing the expression levels of *HMGA1*, *TNFRSF12A*, *CHN1*, *PHLDA2*, and *CDH13*. (**p ≤ 0.01; ***p ≤ 0.001)

**Table S1.** Patient demographics and transplant statistics.

| Characteristic |  | CLAD (n=7) | Control (n=5) | Total (n=12) | P-Value |
| --- | --- | --- | --- | --- | --- |
| Age in years at Transplant, mean (sd) |  | 48.9 (18.5) | 51.4 (13.2) | 49.9 (15.9) | 0.80 |
| Female, n(%) |  | 4 (57.1) | 1 (20.0) | 5 (41.7) | 0.20 |
| Pre-Transplant Diagnosis, n(%) |  |  |  |  |  |
|  | COPD | 2 (28.6) | 2 (40.0) | 4 (33.3) | 0.83 |
|  | Cystic Fibrosis | 1 (14.3) | 1 (20.0) | 2 (16.7) |  |
|  | ILD | 3 (42.9) | 2 (40.0) | 5 (41.7) |  |
|  | Other | 1 (14.3) | 0 | 1 (8.3) |  |
| BAL Characteristics, mean (sd) |  |  |  |  |  |
|  | Volume (mL) | 18.6 (2.7) | 22.4 (10.6) | 20.2 (7.0) | 0.38 |
|  | Cell Count (x10<sup>6</sup>) | 5.3 (6.5) | 3.2 (1.9) | 4.4 (5.1) | 0.51 |
| | Red Blood Cells, cells/μL | 1435.7 (1472.5) | 1517.8 (1711.7) | 1465.6 (1477.1) | 0.93 |
| | White Blood Cells, cells/μL | 884.6 (1020.3) | 541.0 (294.4) | 759.6 (825.0) | 0.54 |
| | Lymphocytes, cells/μL | 6.4 (7.2) | 8.0 (11.5) | 7.0 (8.4) | 0.78 |
| | Neutrophils, cells/μL | 27.6 (27.6) | 25.3 (19.4) | 26.7 (23.9) | 0.89 |
| | Histiocytes, cells/μL | 64.7 (33.3) | 65.8 (31.6) | 65.1 (31.0) | 0.96 |
| | Eosinophils, cells/μL | 1.3 (2.0) | 0 | 0.8 (1.7) | 0.24 |
| Time from Transplant to CLAD onset in days, mean (sd) |  | 1387.3 (787.1) | N/A | N/A | N/A |
| CLAD Phenotype, n(%) |  |  |  |  |  |
|  | BOS | 1 (14.3) | N/A | N/A | N/A |
|  | RAS | 2 (28.6) | N/A | N/A | N/A |
|  | Mixed | 2 (28.6) | N/A | N/A | N/A |
|  | Undefined | 2 (28.6) | N/A | N/A | N/A |

BAL, bronchoalveolar lavage; COPD, chronic obstructive pulmonary disease; ILD, interstitial lung disease

